# Inference of selective sweep parameters through supervised learning

**DOI:** 10.1101/2022.07.19.500702

**Authors:** Ian V. Caldas, Andrew G. Clark, Philipp W. Messer

**Affiliations:** Department of Computational Biology, Cornell University, USA; Department of Molecular Biology and Genetics, Cornell University, USA

## Abstract

A selective sweep occurs when positive selection drives an initially rare allele to high population frequency. In nature, the precise parameters of a sweep are seldom known: How strong was positive selection? Did the sweep involve only a single adaptive allele (hard sweep) or were multiple adaptive alleles at the locus sweeping at the same time (soft sweep)? If the sweep was soft, did these alleles originate from recurrent new mutations (RNM) or from standing genetic variation (SGV)? Here, we present a method based on supervised machine learning to infer such parameters from the patterns of genetic variation observed around a given sweep locus. Our method is trained on sweep data simulated with SLiM, a fast and flexible framework that allows us to generate training data across a wide spectrum of evolutionary scenarios and can be tailored towards the specific population of interest. Inferences are based on summary statistics describing patterns of nucleotide diversity, haplotype structure, and linkage disequilibrium, which are estimated across systematically varying genomic window sizes to capture sweeps across a wide range of selection strengths. We show that our method can accurately infer selection coefficients in the range 0.01 < *s* < 100 and classify sweep types between hard sweeps, RNM soft sweeps, and SGV soft sweeps with accuracy 69 % to 95 % depending on sweep strength. We also show that the method infers the correct sweep types at three empirical loci known to be associated with the recent evolution of pesticide resistance in *Drosophila melanogaster*. Our study demonstrates the power of machine learning for inferring sweep parameters from present-day genotyping samples, opening the door to a better understanding of the modes of adaptive evolution in nature.

**Author summary:** Adaptation often involves the rapid spread of a beneficial genetic variant through the population in a process called a selective sweep. Here, we develop a method based on machine learning that can infer the strength of selection driving such a sweep, and distinguish whether it involved only a single adaptive variant (a so-called hard sweep) or several adaptive variants of independent origin that were simultaneously rising in frequency at the same genomic position (a so-called soft selective sweep). Our machine learning method is trained on simulated data and only requires data sampled from a single population at a single point in time. To address the challenge of simulating realistic datasets for training, we explore the behavior of the method under a variety of testing scenarios, including scenarios where the history of the population of interest was misspecified. Finally, to illustrate the accuracy of our method, we apply it to three known sweep loci that have contributed to the evolution of pesticide resistance in *Drosophila melanogaster*.

## Introduction

When positive selection drives an initially rare adaptive mutation to high population frequency, this leaves the characteristic patterns of a so-called selective sweep in the surrounding genetic variation (Maynard Smith & Haigh, 1974). Over the past 20 years, a variety of summary statistics and computational approaches have been developed for detecting target loci of recent positive selection by searching for sweep signatures (Nielsen et al., 2005; Pavlidis et al., 2013; Sabeti et al., 2007; Tajima, 1989; Vitti et al., 2013). The application of these selection scans has helped us uncover the molecular basis of many examples of recent adaptations, including loci of medical and commercial relevance such as those underlying drug resistance in human or livestock pathogens (Parobek et al., 2016; Redman et al., 2015), and insecticide resistance in crop pests (Anderson et al., 2018; Calla et al., 2021).

The specific patterns a sweep is expected to produce can depend on its parameters and evolutionary history. The strength of positive selection driving the sweep, for example, should determine the size of the genomic region over which a sweep signature can be observed (Kaplan et al., 1989). Furthermore, three different modes of selective sweeps are generally distinguished based on the genealogy of adaptive alleles in a population sample, with each type potentially producing distinct signatures (Hermisson & Pennings, 2005; Pennings & Hermisson, 2006a, 2006b): In the classical “hard” selective sweep, a single adaptive allele arose by mutation and was immediately positively selected. At the adaptive locus, all sampled lineages that carry the adaptive allele should therefore coalesce in a most recent common ancestor that lived after the onset of positive selection. The two other categories constitute so-called “soft” selective sweeps, where the most recent common ancestor of the sampled alleles lived prior to the onset of positive selection. This could be because the adaptive allele already existed in the population at an intermediate frequency before it became adaptive, and multiple distinct lineages from that time were captured in the sample. We then refer to the sweep as a soft sweep from standing genetic variation (SGV). Another possibility is that the adaptive allele arose repeatedly in the population by independent *de novo* mutation events, in which case we refer to the sweep as a soft sweep from recurrent new mutations (RNM).

It is important to recognize that this classification of sweep types is based on the genealogy of adaptive alleles in a given population sample. Consequently, the same adaptive event can result in a sweep that is soft in one sample but hard in another. For example, if the adaptive allele arose several times in the population, a sample capturing lineages from two distinct origins would constitute a soft sweep, while a sample capturing only a single lineage (maybe because that lineage was much more prevalent in the population) would constitute a hard sweep. Generally, we expect hard sweeps to dominate adaptation in mutation-limited scenarios, while soft sweeps should be more common in larger populations where adaptation is not limited by the availability of adaptive mutations due to a high level of standing variation and/or high population-level mutation rates towards adaptive alleles (Hermisson & Pennings, 2017; Messer & Petrov, 2013).

The footprints of soft sweeps can be quite different from those of hard sweeps, and are often more difficult to detect (Berg & Coop, 2015; Ferrer-Admetlla et al., 2014; Peter et al., 2012). Hard sweeps are characterized by a very recent common ancestor of the adaptive allele, with a “star-like” genealogy at the selected site. As a result, their hallmark signatures include a trough in genetic diversity around the adaptive site, the presence of a single long haplotype, and a characteristic skew in the site frequency spectrum (SFS) of linked neutral polymorphisms towards high and low derived allele frequencies (Fay & Wu, 2000; Sabeti et al., 2002). In a soft sweep, by contrast, the longer time to the most recent common ancestor can result in higher levels of genetic diversity being maintained at the sweep locus, with several long adaptive haplotypes possibly present at intermediate population frequencies (Pennings & Hermisson, 2006b; Przeworski et al., 2005). These differences to classical hard sweep signatures should be most pronounced for RNM soft sweeps, whereas SGV soft sweeps can span a range of signatures, from those similar to RNM sweeps to signatures that are essentially indistinguishable from hard sweeps, depending on the specific evolutionary history of the adaptive allele. If the adaptive allele in an SGV sweep was still young at the onset of positive selection (maybe because it was previously deleterious), the resulting sweep signature should be very similar to a hard sweep. Conversely, if the allele was much older and already present on several diverged haplotypes at the onset of positive selection, this should generate a signature more similar to an RNM sweep. The adaptive allele could also have originated multiple times prior to the onset of selection and several alleles of independent origins could have been picked up by selection, which should again produce a signature resembling an RNM sweep.

The fact that sweep mode and parameters can affect sweep signatures raises the possibility that we may be able, in turn, to infer these parameters for a given sweep by analyzing its signatures in a population sample. Such knowledge could provide valuable insights into the nature of adaptive events. Consider, for example, a sweep associated with the evolution of drug resistance in a pathogen such as the malaria parasite *Plasmodium falciparum*. Knowing the strength of selection that drove this sweep could allow us to predict how rapidly the responsible mutations are expected to spread when introduced into a new population, while knowing the mode of the sweep could help us assess whether these mutations can evolve quickly and repeatedly, or whether this was possibly a one-off event.

Indeed, several methods for inferring sweep mode and selection coefficients have recently been developed that can draw such inferences from the polymorphism patterns observed in a single population sample. The popular sweep scans SweepFinder and SweeD (DeGiorgio et al., 2016; Pavlidis et al., 2013) already provide estimates of selection coefficients based on the analysis of the shape of the SFS around a putative sweep locus using maximum likelihood analyses. Other approaches can estimate selection coefficients from the distribution of haplotype frequencies (Messer & Neher, 2012) or inferred ancestral recombination graphs (Hejase et al., 2021; Stern et al., 2019). A shortcoming of these analytical approaches is that they require rigid assumptions such as presuming a panmictic population of constant size and/or fixed sweeps. Even when approaches are robust to violations of their assumptions, it is unclear how well they can be targeted to a specific scenario if there is more information available about the history of the population of interest. Moreover, analytical approaches are based on average sweep signals, but the signal of a given individual sweep is stochastic and may deviate strongly from the analytical expectation. Several methods have further been devised for distinguishing hard from soft sweeps using computational approaches such as Approximate Bayesian Computation (Garud et al., 2015; Peter et al., 2012; Stern et al., 2019), but both analytical and likelihood-free approaches tend to require tuning of *a priori* analysis hyperparameters, including genomic window sizes. This is an important choice because the region over which sweep signatures are expected to extend is approximately inversely proportional to the strength of positive selection that drove the sweep (Kaplan et al., 1989). Thus, by choosing a specific window size, one intrinsically gears a method to a specific selection strength. This is a problem if this selection strength and, therefore, the appropriate window size to capture the sweep, is unknown in advance.

Supervised machine learning provides a new paradigm for evolutionary analyses that has gained increasing attention over the past years (Schrider & Kern, 2018). Under this paradigm, *in silico* polymorphism datasets are simulated and used as training data to fit a statistical model, which is then applied to make inferences from the empirical data. When trained on a distribution of sweep signatures with known evolutionary history, any parameter of a given sweep could in principle be inferred by the model, as long as we can train it with accurate and appropriate simulations. Importantly, due to the flexibility provided by simulations, which can explore large regions of parameter space and be designed to represent any particular organism and locus of interest, supervised machine learning could provide a powerful approach for making sweep inferences for a variety of organisms and scenarios. Indeed, several implementations of supervised machine learning for sweep inferences have already been successfully demonstrated in recent years (Flagel et al., 2019; Kern & Schrider, 2018; Lin et al., 2011; Mughal & DeGiorgio, 2019; Pavlidis et al., 2010; Pybus et al., 2015; Ronen et al., 2013; Schrider & Kern, 2016; Sheehan & Song, 2016; Sugden et al., 2018; Torada et al., 2019; Xue et al., 2021). By their nature of learning by example from diverse training data, these methods are naturally capable of learning patterns across individual sweeps with highly stochastic signatures and across a variety of analysis hyperparameters such as window size.

In this paper, we introduce a novel supervised learning framework that can in principle be trained to infer any evolutionary parameter of a given selective sweep from its observed signature. We present a way to efficiently simulate hard and soft selective sweeps to produce training datasets. We fit convolutional neural networks to estimate sweep mode and selection coefficient of an observed sweep and show an example of extending the approach by comparing models trained on fixed sweeps and ongoing sweeps. Our method achieves good performance on validation datasets, indicating that signatures left by selective sweeps in surrounding neutral polymorphism are informative about their mode and parameters. Finally, we apply the method to previously characterized sweeps associated with pesticide resistance in *Drosophila melanogaster* as an application to a real dataset, confirming that our parameter estimates agree with previous experimentally-derived hypotheses for these loci.

## Methods

Our method for inference of sweep parameters involves four main steps (Fig. 1). First, we generate a large data set of simulated sweep signatures spanning different types and selection coefficients. For each sweep simulation, we calculate a set of haplotypes and SFS-based summary statistics at different genomic locations around the sweep locus, using systematically varying window sizes. The resulting data is split into a training and a validation set. With the former, we train a convolutional neural network (CNN) capable of estimating different sweep parameters of interest. Finally, we apply the trained model to the validation set to evaluate the performance of the model. The method is implemented as a reproducible pipeline where each simulation and analysis parameter can be tuned (Table 1). Parameters can be given a constant value or specified by a probability distribution; the pipeline currently accepts uniform, log-uniform, and integer uniform distributions. In the case of a distribution, a random value is picked from it for every simulation.

**Figure 1:**
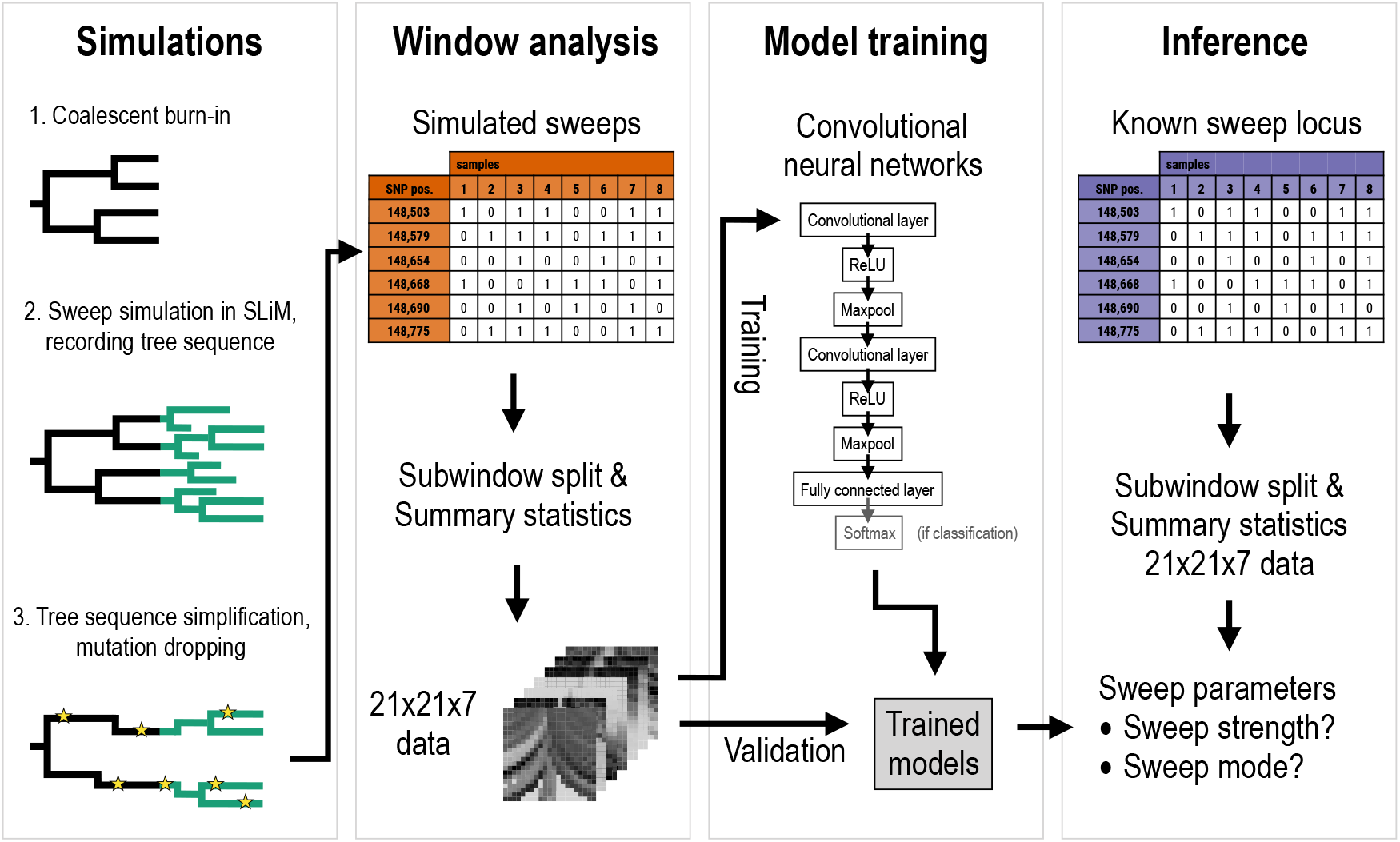
Diagram of inference method.

**Table 1:** Set of tunable simulation parameters in the analysis pipeline. Adaptive mutation rate and frequency at onset of selection are only meaningful for RNM or SGV sweeps, respectively.

| Parameter | Symbol | Training value |
| --- | --- | --- |
| Population size | $N_e$ | 50 000 |
| Number of sampled genomes | $k$ | 205 |
| Total locus size | $L$ | 1000 kb |
| Neutral mutation rate | $\mu$ | $2.25 \times 10^{-8}$ |
| Uniform recombination rate | $r$ | $1.619 \times 10^{-7}$ |
| Sweep site coordinate | $x_{\text{sweep}}$ | 500 kb |
| Selection coefficient | $s$ | log-Uniform(0.01, 100) |
| Dominance coefficient | $h$ | 0.5 |
| Frequency at sampling | $f_{\text{sample}}$ | 1.0 |
| Adaptive mutation rate | $\mu_{\alpha}$ | log-Uniform($5 \times 10^{-8}$, $2.25 \times 10^{-5}$) |
| Frequency at selection onset | $f_0$ | log-Uniform(0.0002, 0.01) |
| Maximum number of restarts | $R_{\text{max}}$ | 1000 |
| Data dimension | $d$ | 21 |
| Smallest subwindow size | $l_{\text{min}}$ | 1 kb |

### Sweep simulations

We model an adaptive site located at position *x*_sweep_ of a genomic region of length *L* base pairs in a diploid population. For simplicity, we assume that this adaptive site is a single nucleotide position. Note that this could be a reasonable approximation also for a larger adaptive target site as long as alleles remain effectively linked across this site during the sweep (Messer & Petrov, 2013). All other sites in the region are assumed to be neutral. We model adaptive alleles with a selection coefficient *s* > 0 such that wild-type individuals are assigned fitness 1, heterozygotes are assigned fitness 1 + *hs*, and homozygotes for the adaptive allele are assigned fitness 1 + *s*.

Our sweep simulations employ a hybrid approach that combines coalescence and forward simulation (Haller et al., 2019). The initial state is a neutral coalescent burn-in generated in msprime (Kelleher et al., 2016), which is saved in the succinct tree sequence format (Kelleher et al., 2018). This tree sequence is then imported into SLiM 3.7 (Haller & Messer, 2019) to simulate the selection phase of the sweep. Importantly, only the trajectory of adaptive alleles and recombination breakpoints occurring at the specified recombination rate *r* are modeled in this phase, but no neutral mutations. The tree sequence is continuously updated by SLiM. After the adaptive allele has reached a desired frequency *f*_sample_, we stop the simulation to obtain a population sample. This is done by importing the resulting tree sequence back into msprime, and then taking *k* random leaf nodes from the tree sequence, corresponding to a sample of genomes of size *k* from the population. The result is a simplified tree sequence representing the entire genealogical history of the sample, on which neutral mutations are then dropped by msprime according to the specified mutation rate μ. Finally, we convert the leaves (samples) into a list of haplotypes in ms format according to the infinite-sites model. This hybrid simulation strategy allows us to leverage the efficiency of coalescent simulations while keeping the flexibility of forward simulations, which can be customized in various aspects of the evolutionary scenario such as demography, genetic architecture, and population life history.

In the selection phase, our simulations can model three types of sweeps: hard sweeps, SGV soft sweeps, and RNM soft sweeps. To model hard sweeps, we introduce a single copy of the adaptive allele with given selection coefficient *s* into a randomly chosen chromosome from the population, and then follow its frequency trajectory. If genetic drift causes the adaptive allele to be lost prior to reaching the desired frequency *f*_sample_, the simulation is reset to the start of the selection phase and the adaptive allele is reintroduced. This is repeated until a sweep of the desired population frequency is obtained.

To model soft sweeps from SGV, we assume that the adaptive allele is initially neutral and drifts to a given population frequency *f*_0_, at which point it first becomes adaptive. This frequency *f*_0_ can be interpreted as a tuning parameter for the “softness” of an SGV sweep, with higher frequencies tending to result in softer sweeps. We simulate this scenario by introducing a single copy of the allele into a randomly chosen chromosome, and then following its frequency trajectory under drift until it is either lost or reaches for the first time the desired target frequency *f*_0_. If lost, we go back to the starting point and reintroduce the allele in a single copy. If the allele successfully reaches frequency *f*_0_, its selection coefficient is then set to the desired value *s* > 0 and its frequency trajectory is further followed until it reaches the frequency *f*_sample_ or is lost. In the latter case, the simulation is reset and repeated from the point where the allele was recorded at frequency f_0_ and positive selection started.

To model soft sweeps from RNM, we assume that new instances of the adaptive allele arise at the selected locus at a specified “adaptive mutation rate” *μ_α_*, such that multiple versions of the adaptive allele from independent mutational origins can contribute to the sweep. The value of *μ_α_* here serves as the tuning parameter for sweep softness, with higher values tending to result in more versions of the adaptive allele segregating in the population and therefore softer sweeps. We simulate this scenario similarly to the hard sweep scenario, except that new instances of the adaptive allele can now continue to arise at the specified rate *μ_α_* while the sweep is progressing. All instances of the adaptive allele are assumed to have the same selection coefficient *s*. However, we allow each chromosome to carry at most one such allele, meaning that if a new adaptive mutation occurs on a chromosome that already carries one, the original adaptive allele will be kept and the new one discarded. The simulation is followed until the combined frequency of adaptive alleles across chromosomes reaches *f*_sample_. Note that similar to our hard sweep simulations, the simulation is restarted from the beginning if the initial first copy of the adaptive allele is lost.

Whenever a simulation is restarted due to the adaptive allele being lost, it is given a new seed for the random number generator in SLiM, but its parameters retain their exact numerical value. This guarantees each set of simulation parameters is given multiple chances to produce an evolutionary trajectory resulting in a selective sweep. For computational purposes, however, a failsafe is implemented where the set of simulation parameters is entirely discarded if the number of simulation restarts exceeds a threshold *R*_max_.

One important thing to keep in mind is that not all sweeps generated by the above SGV and RNM simulation models will indeed be soft. If f_0_ or *μ_α_* are sufficiently small, both models may generate hard sweeps with high probability (Hermisson & Pennings, 2017). To ensure that our machine learning approach is trained with sweeps of the correct type, we reject any simulated sweeps generated under the SGV or RNM models that are not actually soft according to their genealogy at the adaptive site. Specifically, this means we keep only those sweeps generated under the above SGV and RNM procedures where the coalescence time of all sampled adaptive allele copies is indeed older than the onset of positive selection, while sweeps with younger coalescence times are rejected. According to this criterion, under our simulation parameters, there was a probability of rejection of 47 % for RNM simulations and 12 % for SGV simulations.

### Window analysis of sweep signatures

Our machine learning framework is trained on a set of summary statistics; the pipeline currently has implemented 7 of them. Three of these statistics are designed to capture features of the SFS: the total number of SNPs, the average nucleotide heterozygosity *π* (Charlesworth & Charlesworth, 2010), and Tajima’s *D* (Tajima, 1989). The other four are designed to capture features of the haplotype frequency spectrum: the number of distinct haplotypes, and haplotype homozygosity measures *H*_1_, *H*_12_, and *H*_2_/*H*_1_ (Garud et al., 2015). We chose this broad set of statistics because they have already been successfully used in previous approaches and can capture different aspects of polymorphism patterns that may be informative about sweep parameters and type.

These statistics are considered “windowed” in that they require specification of a genomic window over which they are estimated. This choice is to some extent arbitrary and previous approaches have invoked different rationales for specific choices. For example, Garud et al. (2021) estimated H_12_ and H_2_/H_1_ statistics in their study over windows centered on the putative sweep locus, using a window size of 401 SNPs, which corresponds to approximately 10 kb in the *D. melanogaster* population samples they analyzed. The choice of window size intrinsically gears a method to a specific sweep strength, but this could be problematic if the method is intended to be capable of inferring sweep parameters and type over a broad range of selection strengths.

To address this issue, our method adopts a different approach where each statistic is evaluated over a wide range of systematically varying positions and window sizes, and the machine learning model is then trained on all of these data. In particular, we estimate each of the 7 summary statistics over a total of *d* × *d* = *d*^2^ subwindows designed to capture neutral polymorphism at different locations and resolutions around the sweep locus (Fig. 2). The smallest subwindow size is specified in base pairs by the parameter *l*_min_; the largest is the size needed to cover the full genomic region of length *L,* with intermediate sizes scaling logarithmically. Positions of subwindows are chosen such that subwindows of the same size overlap by half their size with each neighbor. The resulting 7*d*^2^ data points for each simulated population sample provide the data representation and input for our machine learning algorithms. Note that in contrast to SNP window sizes, we define our windows by a number of base pairs. This allows window sizes to remain constant over every sweep simulation, no matter their parameters, and lets the heterozygosity in a genomic region be itself part of sweep signature.

**Figure 2:**
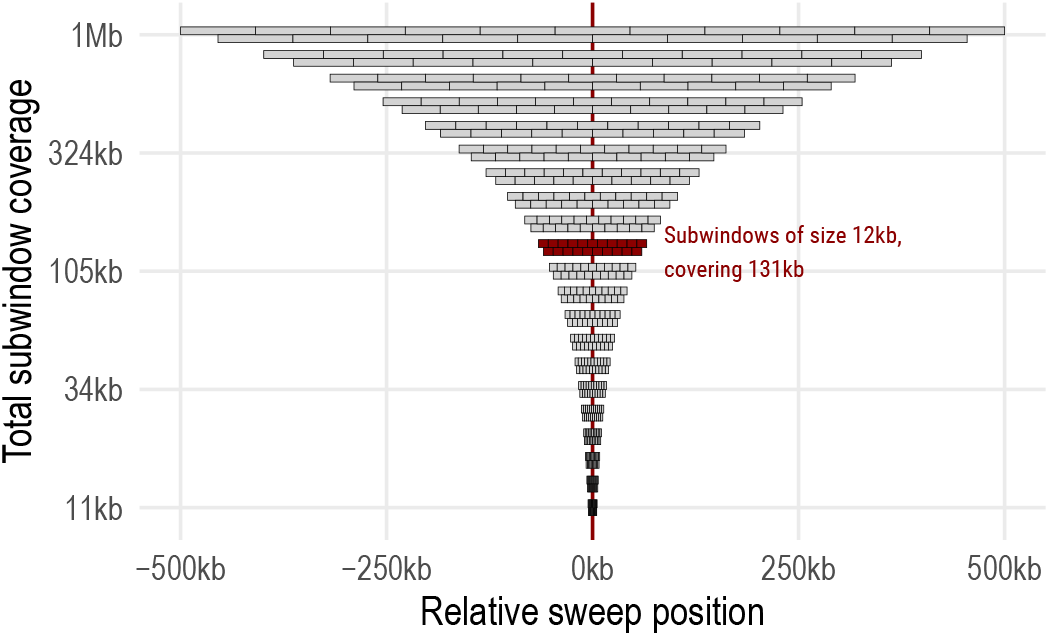
Division of genomic region into subwindows. Base pair values are shown for *d* = 21 dimensions, with minimum subwindow size of *l*_min_ = 1 kb and a total locus size of *L* = 1000 kb. One subwindow size (12 kb) is highlighted in red. Exact window sizes are listed in Table S2.

For the purposes of computational neural network fitting, values are normalized to integers in the range 0 to 255 on a linear scale. Raw values above or below the bounds are then converted into the upper or lower bounds, respectively. Bounds for most remaining statistics were based on biological limits (Table S1). The bounds for Tajima’s *D* were picked as a range of 3 standard deviations above and below its theoretical mean of 0 (Tajima, 1989). For *π* and the total number of SNPs, upper bounds were the highest values observed in subwindows belonging to the empirical control sweeps in *D. melanogaster* data; see below.

### Implementation of machine learning models

For the inference of sweep parameters and mode, our method trains a convolutional neural network (CNN) taking as input the normalized data structure of 7*d*^2^ values. The CNN’s architecture with convolutional filters can take full advantage of the correlation structure between the three dimensions of subwindow location, subwindow size, and summary statistic. The exact network architecure is a tunable parameter in the pipeline and can be freely chosen as long as it accepts as input three-dimensional data of shape 7 × *d* × *d*. The architecture implemented by our pipeline consists of two groups of hidden layers, each composed of a convolutional layer (with 2 × 2 kernels, stride 1, and padding 1), ReLU activation, and Maxpoool regularization (with a 2 ×2 kernel). The first convolutional layer has 128 channels of filters and the second 64. After passing through the second group of hidden layers, data is flattened and passed to a fully connected output layer. For classification models, that output is then passed to an additional Softmax activation layer to generate label probabilitites. The CNN architecture is implemented in PyTorch (Paszke et al., 2019). To avoid overfitting and shorten training time, our pipeline employs the 1cycle learning policy of Smith (2018), as implemented in the fastai v2 library (https://github.com/fastai/fastai).

### Application to positive controls in *D. melanogaster*

To test our method on empirical data, we used three previously studied selective sweeps in *D. melanogaster* genes associated with the evolution of pesticide resistance: *Ace* (FlyBase ID FBgn0000024); *CHKov1* (FlyBase ID FBgn0045761); and *Cyp6g1* (FlyBase ID FBgn0025454) as positive controls. Our analyses were performed on version 2 of the Drosophila Genetic Reference Panel (DGRP2; Huang et al., 2014; Mackay et al., 2012), which we filtered for biallelic SNP sites with at most 15 % missing data. The data was then imputed with Beagle 5.1 (Browning et al., 2018). From the imputed SNP dataset we extracted subwindows centered around the three sweep loci of interest. The SNP coordinates of the three resistance loci in *Ace* in the DGRP2 are 3R:9 069 054, 3R:9 069 408, and 3R:9 069 721; we used the middle SNP at position 9 069 408 as the center of the *Ace* window. The *CHKov1* window was centered at 2R:21150000, roughly the middle of the gene as recorded in FlyBase. The *Cyp6g1* window was centered at 2R:8 072 884, the insertion point of the *Accord* transposable element that is common to all adaptive alleles at this locus (Battlay et al., 2018). All three control loci represent partial sweeps in the DGRP2 dataset: 78 out of 205 (38 %) of lines have at least one alternate allele at any of the *Ace* resistance loci; 139 lines (67.8%) were found by PCR to have the resistance insertion at the *CHKov1* locus (Magwire et al., 2011); and 155 lines (75.6 %) have a resistant allele at *Cyp6g1* as indicated by the alternate allele at the *Accord* insertion point. For the analysis of genome-wide patterns, we studied 1-Mbp-long windows across the 2L, 2R, 3L and 3R chromosomes at 200 kb steps. Empirical SNP genotypes at the extracted windows were converted to ms format and processed into 21 × 21 × 7 data as described above, using the same statistic normalization bounds.

### Software availability

The code used for simulation and inference in this paper is available at https://github.com/ianvcaldas/drosophila-sweeps, together with instructions on how to adapt the method to new datasets.

## Results

In principle, our machine learning framework can be trained to infer any sweep parameter in any evolutionary scenario that can be appropriately simulated. Below, we first illustrate this for an application of our method to infer selection coefficient and sweep type in a simple population model broadly inspired by *Drosophila melanogaster*. Using this model, we evaluate the method’s performance under different training procedures and its robustness to confounding factors such as misspecified demography or recombination rate, which will provide insights into the internal representation of sweep parameters in our method. We then demonstrate an extension of the method to partial sweeps. Finally, we evaluate the performance at positive controls provided by three known recent selective sweeps in *D. melanogaster*.

### Basic model training and validation

As an initial demonstration of our framework we trained it for inferring selection coefficient and sweep type in a basic model of a diploid panmictic population of constant size. Our choice of parameters for this model was broadly inspired by a natural population of *D. melanogaster* from North Carolina, described in the DGRP2 data set, which we rescaled for computational efficiency to an effective population size of *N_e_* = 50 000 (Haller et al., 2019). The mutation rate was chosen such that the average nucleotide heterozygosity in our model (under neutrality) equaled the empirical genome-wide estimate of *π* = 0.004518 from the DGRP2 data, yielding a value of *μ* = *π*/4*N_e_* = 2.25 × 10^-8^. The recombination rate was chosen such that the ratio of *μ*/*r* in our model equals a previously derived estimate for *D. melanogaster* (Arguello et al., 2019), yielding *r* = 1.619 × 10^-7^. When comparing these values with an empirical estimate of the actual nucleotide mutation rate of *μ*′ = 2.8 × 10^-9^ in *D. melanogaster* (Keightley et al., 2014), this yields a rescaling factor of *μ*/*μ*′ ≈ 8.03. In other words, one generation in our simulations should correspond to approximately eight generations in the real-world population.

The simulated genomic region is of size *L* = 10^6^ base pairs, and we assume a sample size of *k* = 205 chromosomes drawn randomly from the population, equaling the number of inbred lines in the DGRP2. To confirm that this model indeed provides a reasonable approximation for genome-wide polymorphism patterns in the DGRP2, we performed 500 neutral coalescence simulations under the chosen parameters and compared the simulated site-frequency spectra to the empirical spectrum observed in DGRP2 data, showing excellent agreement (Fig. S1).

To generate selective sweep data for model training, we simulated hard sweeps and soft sweeps from RNM and SGV with randomly drawn selection coefficients and softness parameters. Sweep location was set at the center of the simulation region, with sweep coordinate at base pair position *x*_sweep_ = 500000. We further assume that the population samples are taken in the generation where the combined frequency of all adaptive alleles reaches *f*_sample_ = 1.0, i.e., the moment the sweep reaches fixation in the population. For all three types of sweeps, the value of *s* for each simulated sweep was drawn from a log-uniform distribution with bounds 0.01 < *s* < 100. This corresponds to sweep signatures extending over approximately 10 kb to 10 000 kb in our model, thus spanning a wide range of possible sweep signatures in *D. melanogaster*. Simulations of hard sweeps at the extremes of this parameter range illustrate how our window analysis with systematically varying window sizes can capture signatures across this full range of selection coefficients (Fig. S2).

For RNM sweeps, we drew the value of the adaptive mutation rate from a log-uniform distribution with bounds 5 ×10^-8^< *μ_α_* < 2.5 × 10^-5^. This corresponds to a population level adaptive mutation rate *θ_α_* = 4*N_e_μ_α_* between 0.01 and 5, which covers a broad range of softness levels, from hard sweeps to very soft sweeps with many independently originated adaptive alleles captured in the sample (Hermisson & Pennings, 2017). Note, however, that only true soft sweeps (see Methods) were kept and labelled as RNM soft sweeps, while simulations resulting in hard sweeps were discarded. The final set of RNM soft sweeps generated by this procedure contained 2 to 26 (median 4) independently originated adaptive alleles per sample.

For SGV sweeps, the starting frequency *f_0_* at which a previously neutral allele becomes adaptive was drawn from a log-uniform distribution with bounds 2/(2*N_e_*) < *f_0_* < 0.01. This means the number of chromosomes in the population carrying an adaptive allele at the onset of selection ranged from 2 to 1000. Again, only true soft sweeps were kept and labelled as SGV soft sweeps. In the final set of SGV soft sweeps generated by this procedure, the number of different lineages present at onset of positive selection that were captured in the sample ranged from 2 to 188 (median 24).

Overall, we generated a data set of 15 000 sweeps. Our training dataset consisted of 4000 sweeps from each of the three different modes (hard, RNM soft, and SGV soft), and our validation dataset of 1000 sweeps from each mode. The population parameters of the basic model are summarized in Table 1. To calculate windowed summary statistics, we chose a number of subwindow sizes and number of subwindow positions per size of *d* = 21, resulting in 21 × 21 × 7 = 3087 summary statistic values per simulation. The smallest subwindow size was set at *l*_min_ = 1 kb, with larger sizes increasing exponentially as described in the Methods.

We chose selection coefficient and sweep mode as the main evolutionary parameters of interest. Estimation of selection coefficient was implemented as a regression model to determine the base-10 logarithm of the selection coefficient *s* of a complete sweep in our basic model. Estimation of sweep mode was implemented as a three-way classification: given a sweep signature, the method should tell whether it comes from a hard, RNM, or SGV sweep. For each of these two applications, we trained a separate CNN.

To pick the length of training, we performed 10 training replicates of every model with a different training and validation split of the total data. We checked their learning curves against a variety of early stopping criteria designed to avoid overfitting (Prechelt, 2012), and the final training period was chosen as the one producing the lowest stable value of the loss function on the validation dataset (Fig. S3). Further training would cause an even lower decrease of training loss but a gradual increase in validation loss, indicating overfitting. Each model was thus trained for 50 epochs, each epoch being a full pass across the training dataset in batches of size 64.

### Performance evaluation

We first checked the performance of our CNN trained for estimating selection coefficient on the validation dataset. Fig. 3A shows that this CNN estimates the selection coefficient in an unbiased way over all four orders of magnitude of s, for all three selective sweep modes. The regression model of s achieved a validation root mean squared error (RMSE) of 0.11. Since the model operates on a log_10_ (*s*) scale, we define the “mean relative error” (MRE) of the inferences to be (|*s*_true_ – *s*_inferred_|)/*s*_true_, measuring the average amount by which inferences are off compared to the true value. Overall, selection coefficient inference achieves a MRE of 16.9 %. Sweeps from SGV carry less signal about selection strength if compared to hard or RNM sweeps: the MRE for hard sweeps is 14.7%, for RNM sweeps 13.8 %, and for SGV sweeps 22.2 %. There is more uncertainty with increasing selection coefficient (Fig. 4).

**Figure 3:**
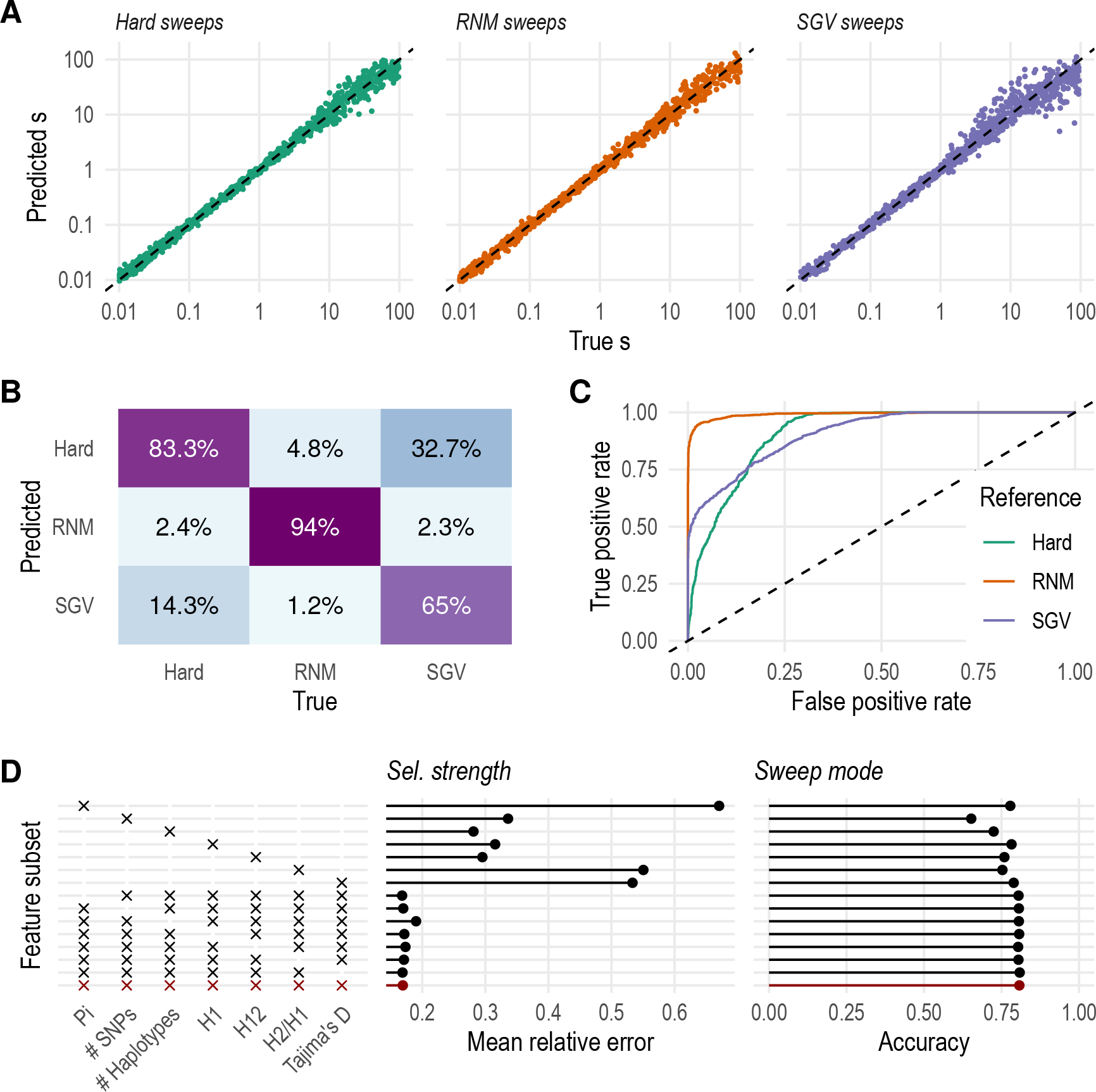
Validation of machine learning models to infer selection coefficient and sweep mode. (A) True versus inferred log_10_ (*s*) for the three sweep modes. (B) Confusion matrix of sweep mode inference, with percentages given across columns. (C) ROC curves for sweep mode inference. Each curve designates a one-vs.-all comparison between a reference mode and the other two modes combined. (D) Inference of selection coefficient and sweep mode by subset of summary statistic. Each row is a separate scenario with only the statistics marked with an “x” on the left panel included in training. The baseline scenario with all statistics is highlighted in red.

**Figure 4:**
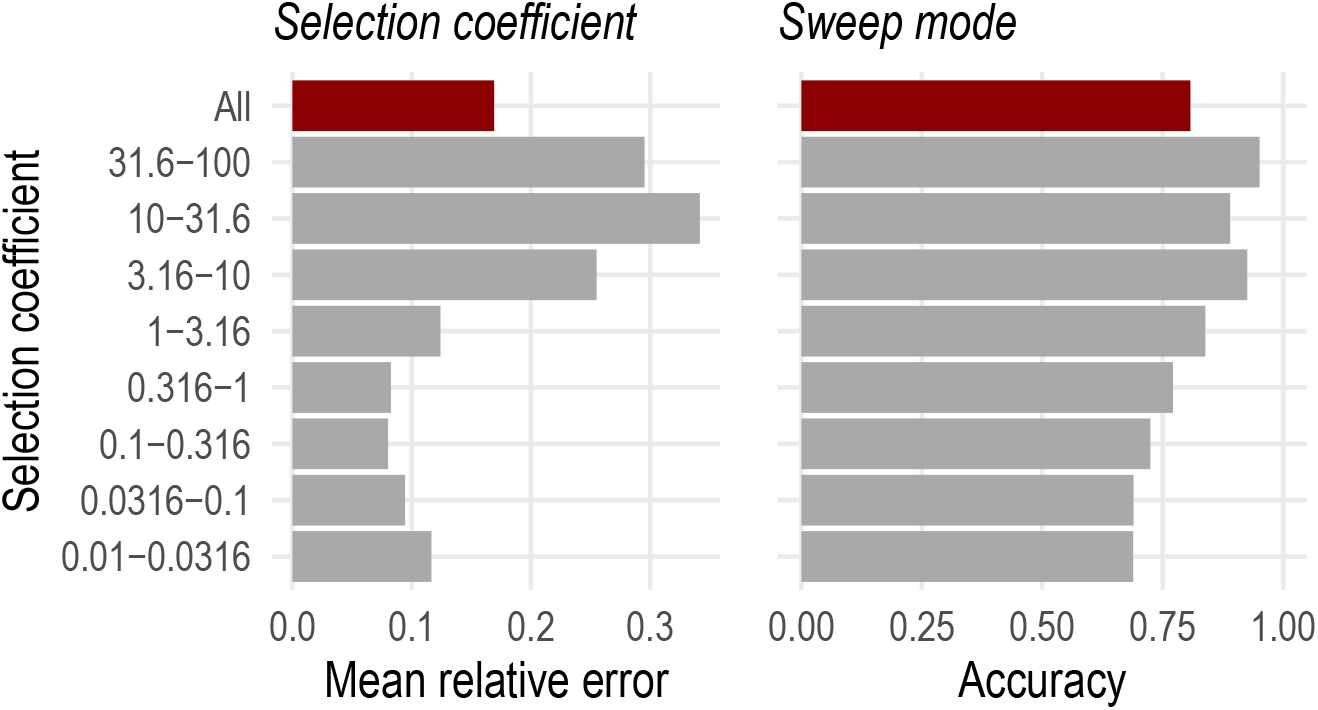
Validation of machine learning models to infer selection coefficient and sweep mode, split by different bins of the selection coefficient s. Each bin contains approximately 380 simulations.

Our CNN trained for sweep mode classification achieves an accuracy of 80.8 % on the validation dataset (Fig. 3B). The model has an average area under the receiver operating characteristic (ROC) curve of 0.936 (Fig. 3C). Overall, our classification performs substantially better than a random guess, which would have 33 % accuracy and area under ROC curve of 0.5. In contrast to selection coefficient, performance of classifying sweep mode increases with sweep strength (Fig. 4). Most mistakes in identifying sweep mode are made when true SGV sweeps are erroneously classified as hard. This is consistent with the fact that some SGV sweeps from low *f*_0_ can have signatures almost indistinguishable from those of hard sweeps.

To aid in interpreting the contributions of the individual summary statistics to the performance of our models, we conducted a feature analysis. For each of the seven statistics used to summarize the sweep signal, we re-trained our models with the same training and validation datasets, but with input modified as to either contain only the statistic of interest (21 × 21 × 1 values per simulation) or all but the statistic of interest (21 × 21 × 6 values per simulation). CNNs were adapted to accept the different input dimensions. Results are shown in Fig. 3D. There is no single statistic that carries the most signal for selection coefficient or sweep mode, and any statistic can be removed from the analysis without great loss of performance. Individual statistics have a more variable distribution of performance.

### Gradient-boosted trees perform comparably to deep learning

Our use of CNNs was motivated by their innate capacity to incorporate correlations across data dimensions. We also trained alternative models to see if CNNs represented a big improvement over an approach that does not involve deep learning. In particular, we used the same datasets to train gradient-boosted tree models (Hastie et al., 2009). Hyperparameters of the models were chosen according to a description of gradient-boosted trees previously shown to work well for many different bioinformatic scenarios and applications (Olson et al., 2017). Validation performances are shown in Table S3. The neural networks have improved performance over the tree-based model, but the difference between the approaches was small.

### Additional binary classification models

The same supervised learning framework can be used for applications where the research question is narrowed. To illustrate this, we trained two additional binary classification models. The first model was trained to distinguish between hard and soft sweeps of any kind. The second model was trained to detect whether a given a soft sweep originated from recurrent *de novo* mutations or standing genetic variation. For the first model, we modified datasets such that RNM and SGV sweeps were given the same label. For the second model, only soft sweeps were included in the training dataset, as hard sweeps are irrelevant to the question. Datasets were balanced such that each label was equally represented, and hyperparameter training proceeded as described previously, with 50 epochs of training (Fig. S3C, D).

The two additional binary classification models performed with high accuracy on validation data (Fig. S4). The first model was able to distinguish between a hard sweep or a soft sweep of any mode with accuracy of 82.9 % and area under ROC curve of 0.905. The second model was able to detect whether a given soft sweep came from recurrent de novo mutations or from standing genetic variation with accuracy 96.1 % and area under ROC curve 0.993. This high performance corroborates that these two different modes of soft sweeps indeed leave distinct genomic signatures from each other. Overall, these results demonstrate that the best option of what simulated datasets to use and what sweep parameter to infer depend on the research question under consideration.

### Misspecification of recombination rate and effective population size

The exact evolutionary parameters of a study population are usually unknown. This raises the question of how sensitive our method will be to misspecification of the parameters used for model training. To examine this question, we first studied the performance of our method under a scenario of equilibrium demography where only the recombination rate was misspecified. This is especially salient because recombination rates vary along the genomes of most organisms, and complete knowledge of the local recombination landscape is not often available. In particular, we applied the models to validation datasets simulated with a recombination rate that was three times higher or lower than the value used for the model training (Table 2). Figure 5A (left two panels) shows that when the actual recombination rate is lower than what the model was trained on (i.e., *r* was overestimated in the training model), sweep coefficient is also systematically overestimated by our method, and vice versa.

**Table 2:**
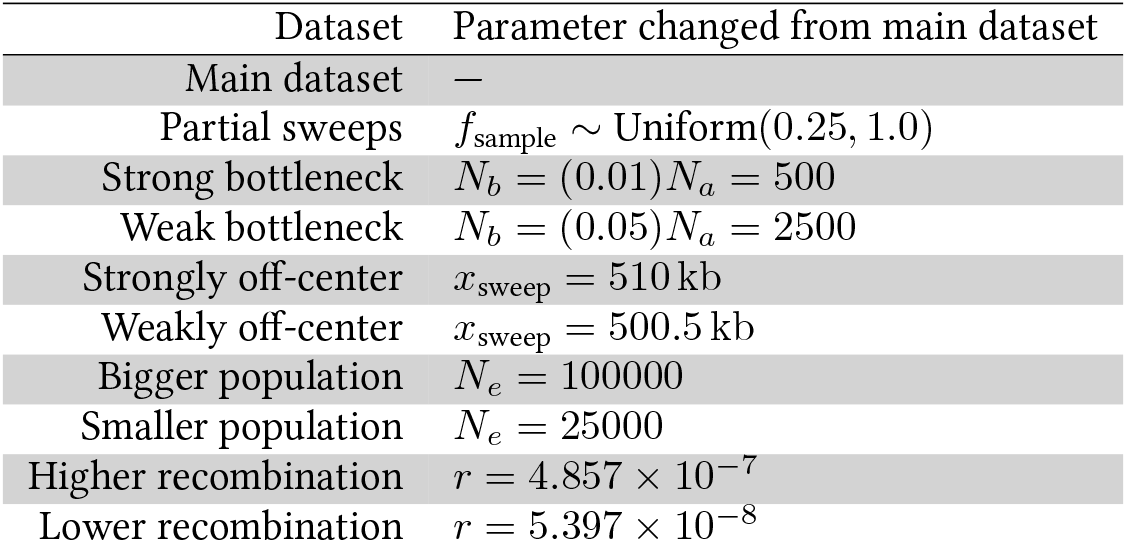
Simulation parameters for all datasets, indicating parameter changes from values in Table 1. The partial sweeps dataset is used for training and validation as well as for assessing the robustness of the model trained on fixed sweeps. *N_a_* refers to the pre-bottleneck population size, while *N_b_* refers to the population size during the bottleneck (see text for full description of the bottleneck simulation strategy).

| Dataset | Parameter changed from main dataset |
| --- | --- |
| Main dataset | — |
| Partial sweeps | $f_{\text{sample}} \sim \text{Uniform}(0.25, 1.0)$ |
| Strong bottleneck | $N_b = (0.01)N_a = 500$ |
| Weak bottleneck | $N_b = (0.05)N_a = 2500$ |
| Strongly off-center | $x_{\text{sweep}} = 510 \text{ kb}$ |
| Weakly off-center | $x_{\text{sweep}} = 500.5 \text{ kb}$ |
| Bigger population | $N_e = 100000$ |
| Smaller population | $N_e = 25000$ |
| Higher recombination | $r = 4.857 \times 10^{-7}$ |
| Lower recombination | $r = 5.397 \times 10^{-8}$ |

**Figure 5:**
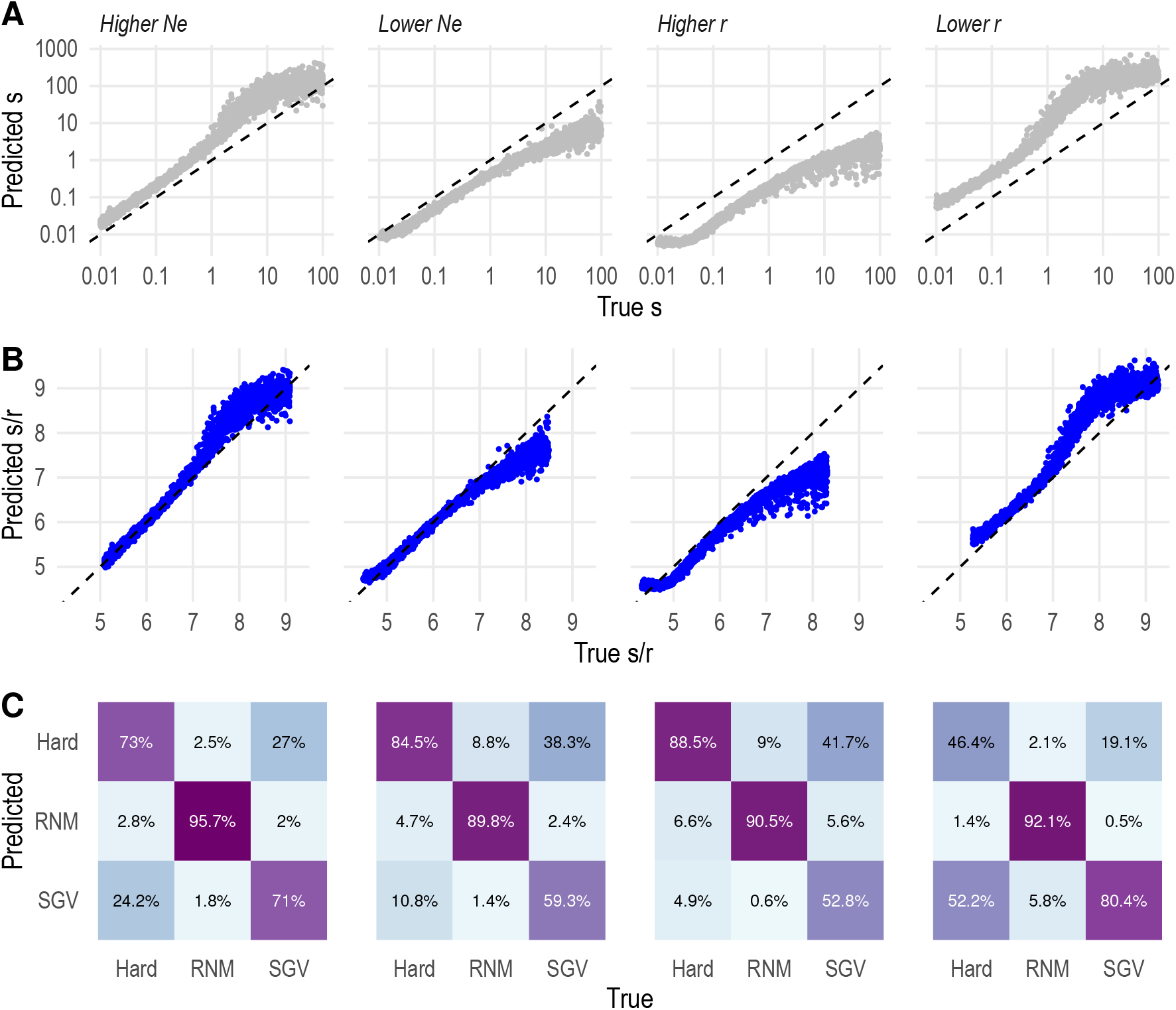
Performance of machine learning models when applied to datasets with values of *r* and *N_e_* different to the training ones. In panel (B), inferred and true *s*/*r* refer to *s*_inferred_/*r*_training_ and *s*_true_/*r*_true_, respectively.

These results are consistent with an interpretation where the neural network for selection coefficient estimation learned to use information on sweep size for its inferences. The expected size of the genomic region over which a sweep signatures extends should be roughly proportional to the inverse of the product of the recombination rate r and the expected sweep duration *τ*, defined as the average number of generations it takes a positively selected mutation destined to fixation to proceed from its initial emergence to fixation in the population (Kaplan et al., 1989). Using the theoretical approximation that *τ* ~ 2ln(2*N_e_s*)/*s* for a codominant mutation of selection coefficient *s* (Desai & Fisher, 2007), this yields an expected sweep size on the order of ~ *s*/[2*r* ln(2*N_e_s*)]. Neglecting the logarithmic dependence on *N_e_s* for now, sweep size should therefore be roughly proportional to the ratio *s*/*r*. Consequently, if the model is indeed trying to fit sweep size, it should compensate for an overestimation of *r* in the training data by also overestimating the inferred s to obtain a sweep of similar size as observed in the actual data, and vice versa.

The behavior of our method when other evolutionary parameters are misspecified further corroborates this interpretation. For example, we studied a scenario where *N_e_* is misspecified in training while Θ = 4*N_e_μ* and the ratio *μ*/*r* are set to their correct values. Such a scenario might be motivated by a study system for which we have an estimate of the level of nucleotide heterozygosity that allows us to infer Θ, as well as an estimate of the relative strength of mutation versus recombination, but we do not know the precise values of *N_e_*, *μ*, and *r*. In that case, one could set one of the parameters, say *N_e_*, to some chosen value, and infer the values of the other two using the two given relations.

Figure 5A (right two panels) shows the results for two examples of such a scenario, where *N_e_* in the validation data was set to a value either two times higher or lower than the value used for model training, while Θ and *μ*/*r* were at their correct values. Here, selection coefficient is underestimated when *N_e_* was overestimated in training, and vice versa. This is again consistent with the above interpretation, because when *N_e_* is overestimated, *μ* will be underestimated, given that Θ = 4*N_e_μ* is kept constant. Since *μ*/*r* is also kept constant, this means *r* will be underestimated in training as well, which the method should compensated for by by an underestimation of the inferred s to obtain a sweep of the size observed in the validation data.

To more directly test our interpretation that the method captures information about sweep size for its selection coefficient inferences, we calculated for each of the above datasets with misspecified training parameters the ratios *s*_inferred_/*r*_training_ and *s*_validation_/*r*_validation_. If the method indeed relies on sweep size for its inferences, these two ratios should be similar, given that sweep size should scale roughly with *s*/*r*. Fig. 5B confirms that this is indeed the case, at least until selection becomes very strong, which makes sense given that the *s*/*r* scaling is expected to break down for large *s*. We conclude moreover that our model is not attempting to simply fit the observed value of the product 2*N_e_s*, a measure of the “effective” coefficient of selection often used in the context of deleterious mutations. If that would be the case, our method would be expected to underestimate s when *N_e_* is overestimated in training, and vice versa, the exact opposite of what is actually observed.

Importantly, the overall accuracy of our method for sweep mode classification was not severely affected by any of the datasets with misspecified training parameters (Fig. 5C). The method performed at a similar overall accuracy of 73.9 % for underestimated r and 75.6 % for overestimated r. For the scenarios where *N_e_* was misspecified, the method performed with accuracy of 77.3 % for underestimated *N_e_* and 80.5 % for overestimated.

### Sweep mode inference is robust to demography misspecification

Demographic events such as population bottlenecks can distort the signatures of selective sweeps (Crisci et al., 2012; Simonsen et al., 1995; Thornton et al., 2007). This could lead to errors in the inferences of sweep parameters if the model is not trained under the correct demographic history. To test the robustness of our method under such demographic misspecification, we applied the models trained on equilibrium demography to sweeps simulated in populations that had undergone a bottleneck. We specifically tested two scenarios where population size was reduced for 100 generations to either 5 % or 1 % of its original value (Table 2).

The onset of the bottleneck in each given simulation run was chosen independently of the start time of the sweep, with each set to happen at a random generation in the range 1 to 2500 after burn-in. Population samples were again taken in the generation where the sweep reached fixation. That way, a sweep could in principle start before, during, or after the bottleneck. However, we discarded those simulations in which the sweep had already fixed prior to bottleneck onset. All sweeps whose trajectories intersected the bottleneck were kept, as were those where the sweep had started after the bottleneck to represent scenarios where a sweep happens in a population recovering from a past reduction in size.

Fig. 6 shows that the presence of a bottleneck can cause overestimation of selection coefficient in our models trained on constant demography, with the effect being larger for the stronger bottleneck scenarios. This overestimation is most pronounced for weaker sweeps with trajectories that overlap with the bottleneck for a substantial period and then ultimately become fixed during the bottleneck; those sweeps are marked in red in Fig. 6A.

**Figure 6:**
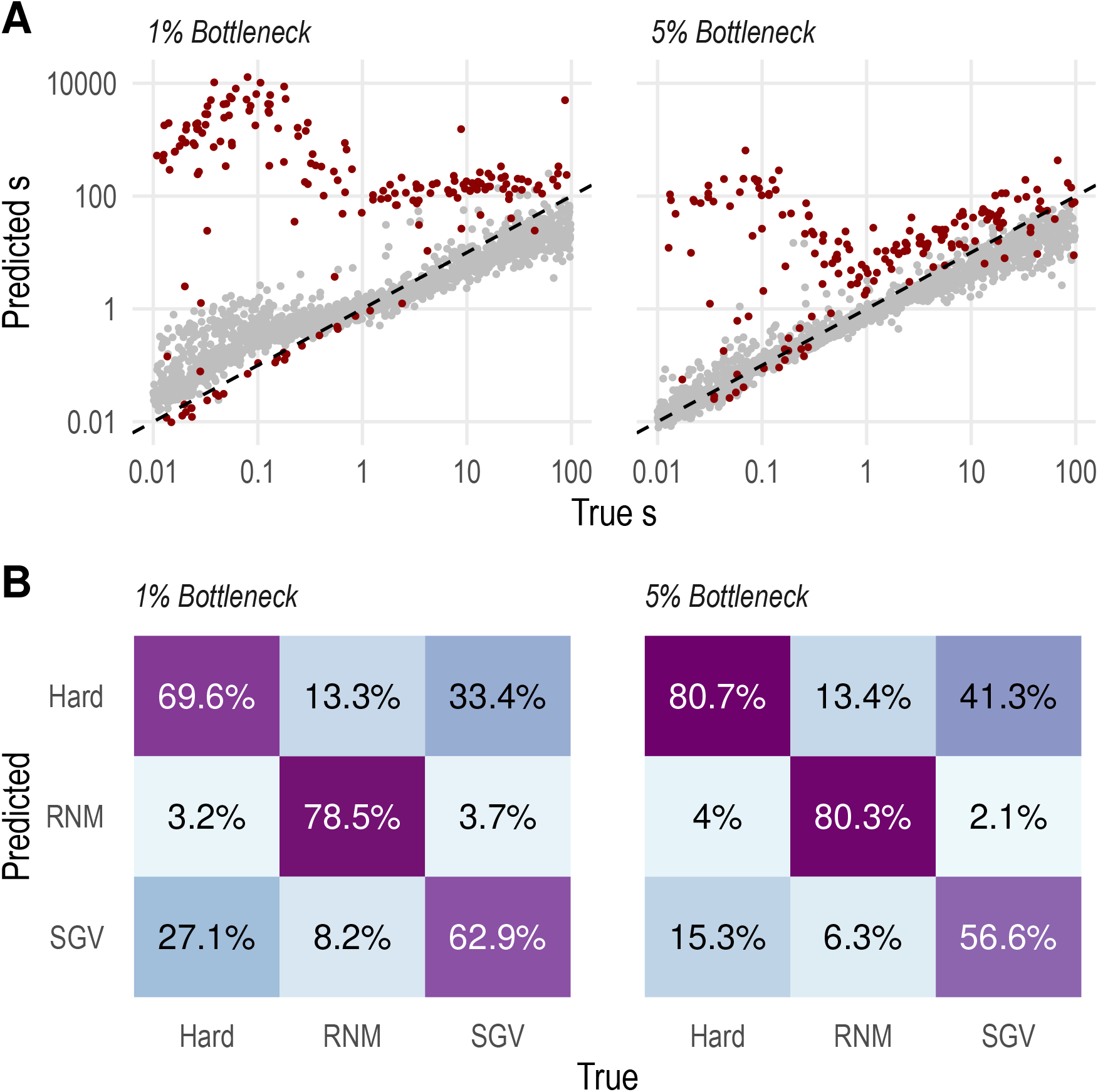
Performance of machine learning models when applied to two datasets with historical bottlenecks. In panel (A), sweeps marked in red have reached fixation during the bottleneck.

This behavior is again consistent with the above interpretation that estimation of selection coefficient is based to some extent on sweep size. Consider, for example, a sweep that would fall entirely inside the bottleneck period (i.e., one that starts and fixes during bottleneck). During its entire “lifetime”, such a sweep would therefore experience the much smaller bottleneck *N_e_*. This would result in a much shorter expected fixation time, and thus a larger sweep size, as compared to a sweep of the same selection coefficient in a population of the original size. Thus, we would expect that our method trained on a model with the constant, larger *N_e_* would overestimate selection coefficient. The relative increase in sweep size, and thus the expected degree of overestimation of *s*, is larger for smaller selection coefficients, consistent with our observations in Fig. 6.

Classification of sweep mode likewise loses power under a bottleneck, performing with an accuracy of 71.5% or 70% for the weaker or stronger bottleneck, respectively. Our model tended to misclassify soft SGV sweeps as hard, and vice-versa, as they did under equilibrium demography. While estimates of selection coefficient can be misled in a predictable direction by the presence of a bottleneck not accounted for in training, the genomic signature of sweep mode is more robust to a temporary reduction in population size, reinforcing the hypothesis that information about different selective sweep parameters is contained in different aspects of the patterns of neutral polymorphism.

### Robustness to sweep mislocalization

Our method assumes we know the precise location of the sweep, but that information might not be so clear in reality. To test the robustness of our method to mislocalization of the sweep, we applied the trained model to two datasets where sweeps were located 0.5 kb and 10 kb away from the exact center of the analysis window (Fig. S5). Selection strength inference was very robust to mislocalization, with MRE of 17.26 % and 17.77 % for mislocalizations of 0.5 kb and 10 kb, respectively. Accuracy of sweep mode inference was very robust to a small mislocalization of 0.5 kb, remaining at 80.6 %, but suffered greatly when the sweep was mislocalized by 10 kb, dropping to 55 %.

### Models trained on fixed sweeps perform poorly on partial sweeps

The training and validation data we have used to this point modeled fixed sweeps. However, partial sweeps could be very common in nature (Pritchard et al., 2010; Ralph & Coop, 2010), and it may not always be straightforward to determine whether a given sweep is fixed or partial. To test how the models trained on fixed sweeps behave when applied to partial sweeps, we generated a validation dataset of sweeps that were sampled when the adaptive allele first reached a given population frequency *f*, drawn from a uniform distribution in the range 0.25 to 1.0 (Table 2). Figure 7 shows that such partial sweeps can confound our method quite substantially. In particular, selection coefficients are underestimated, with the effect being most pronounced for partial sweeps of lower frequencies. Sweep mode is always classified as a soft sweep from RNM, independent of the true mode, leading to essentially random performance. Both effects may be due to the fact that in a partial sweep there are still neutral haplotypes segregating at the sweep locus, resulting in higher levels of genetic variation as compared to a fixed sweep. This could bias our method towards inferring sweep scenarios from the training data of fixed sweeps that maintained the highest levels of diversity, which should be RNM soft sweeps with weak selection. Our results confirm the previous findings of Xue et al. (2021) that models trained on fixed sweeps are not robust when applied to partial sweeps.

**Figure 7:**
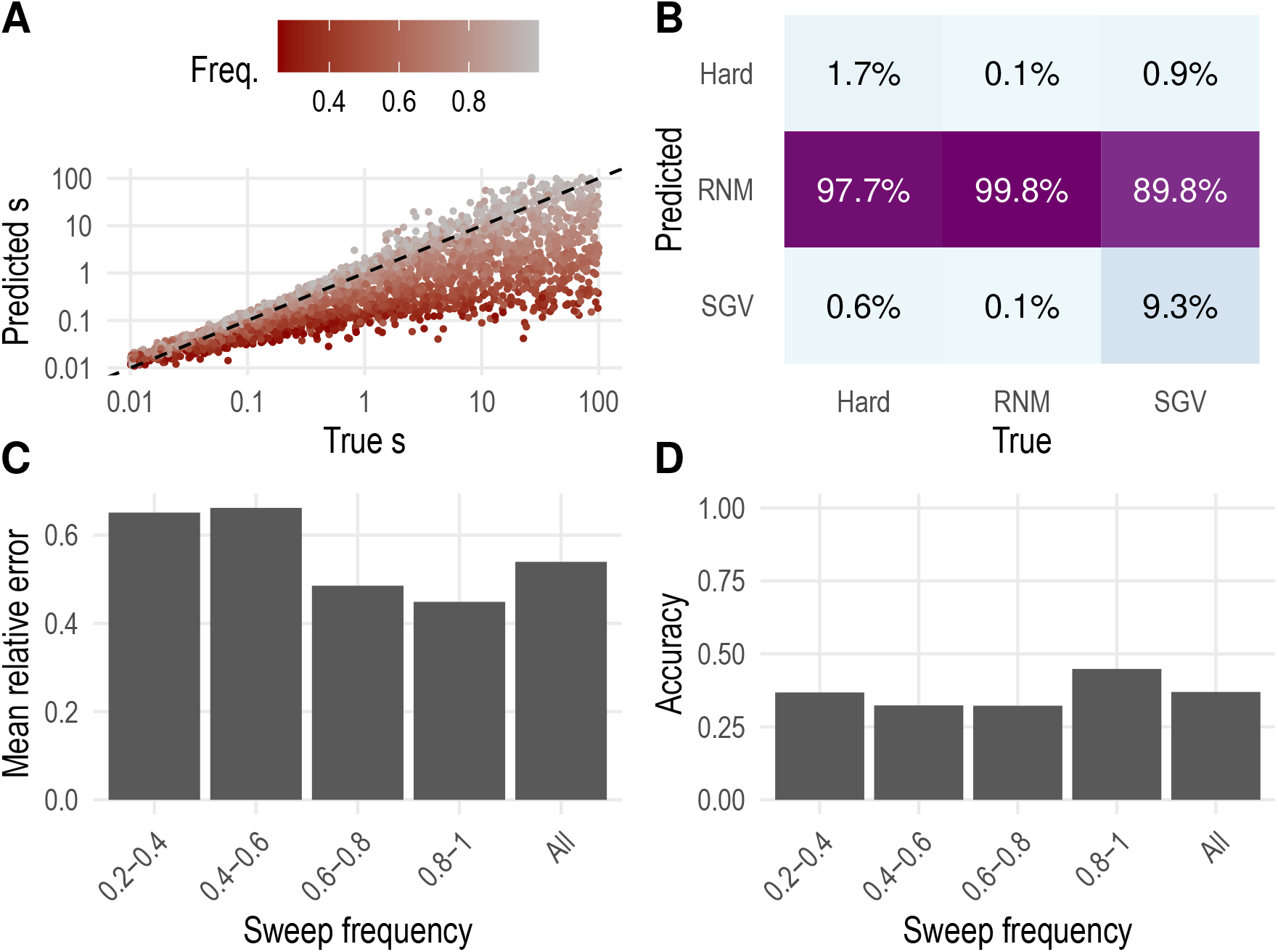
Performance of machine learning models when applied to a dataset of partial sweeps. Sweep frequencies at time of sampling were distributed uniformly in the range 0.25 to 1.0.

### Extending the model to partial sweeps

Given the observation that a model trained on fixed sweeps performs poorly when applied to partial sweeps, we wanted to test whether explicitly including partial sweeps in model training allows the method to regain its power. For this re-training, we used the same dataset as in the previous section, where sweeps were sampled when the adaptive allele first reaches a given population frequency *f*, drawn from a uniform distribution in the range 0.25 to 1.0 (Table 2). We again split this training dataset into 4000 sweeps of each sweep type for training, and 1000 sweeps of each type for validation. Training proceeded in the same way as for the original dataset, for 50 epochs. No overfitting was observed (Fig. S6).

Figure 8 shows that this re-trained model achieved almost the same accuracy for selection inference as the original model that was trained and validated exclusively on fixed sweeps. Importantly, inference of selection coefficients in this new model was unbiased across the whole range of selection strengths tested. Classification of sweep mode performed at an accuracy of 74.6 %, which is only somewhat lower than the 80.8 % of the original model. Overall, these results suggest that it is critical to include partial sweeps in model training whenever the method is applied to sweeps that may not be fixed in the population.

**Figure 8:**
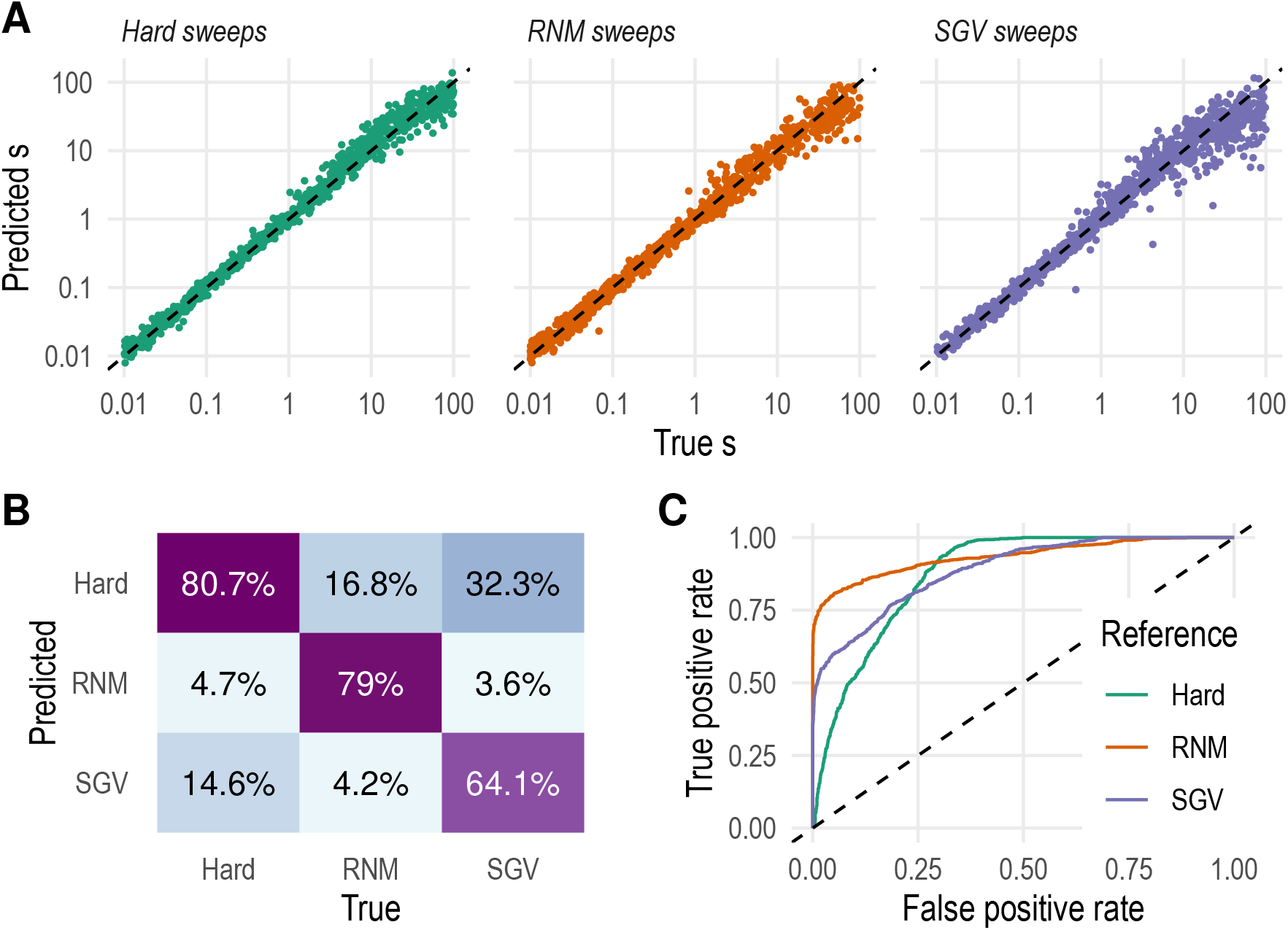
Validation performance of model trained on partial sweeps. (A) True versus inferred log_10_ (*s*) for the three sweep modes. (B) Confusion matrix of sweep mode inference, with given across columns. (C) ROC curves for sweep mode inference. Each curve designates a one-vs.-all comparison between a reference mode and the other two modes combined.

### Performance at known sweep events in *D. melanogaster*

To assess the performance of our method on real-world data, we applied it to three positive control loci in *D. melanogaster* where recent adaptations of known biological mechanisms have left distinct sweep signatures in the DGRP data (Figure S7). The first locus is the gene *Ace*. Here, several point mutations that confer resistance to a variety of pesticides have independently evolved and recently spread through the population (Duneau et al., 2018; Fournier et al., 1993; Karasov et al., 2010). This locus should therefore represent a soft sweep from recurrent *de novo* mutation. The second locus is the gene *CHKov1*, where the recent sweep of a transposable element underlies the evolution of resistance to organophosphates (Aminetzach et al., 2005). Prior to its spread, this transposable element was already segregating at low frequencies in ancestral African populations (Magwire et al., 2011), presumably making this a soft sweep from standing genetic variation. The third locus is the gene *Cyp6g1,* at which a series of nested transposable element insertions followed by a duplication are associated with the recent evolution of resistance to DDT and other pesticides (Daborn et al., 2001; Schmidt et al., 2010). Since multiple adaptive alleles have swept at this locus, it fulfills the definition of a soft sweep. However, given the complex genetic structure of this adaptive event, it is not immediately clear whether it more closely resembles our simulated RNM or SGV soft sweep scenarios.

Our inference models were already trained on parameters chosen to resemble the DGRP2 data (although with a rescaled effective population size of *N_e_* = 50 000 corresponding to a rescaling factor of approximately 8). Thus, we directly applied these models to the three control loci, using as input a window centered at each sweep’s location (see Methods). Since all three sweeps are partial, with adaptive alleles segregating between 30 % and 76 % in the DGRP2, we used the models trained on partial sweeps for these inferences. We trained 10 models with the same training and validation datasets for each inference target to capture the distribution of inference uncertainty.

The fact that inferences are based on a rescaled model has important implications for the interpretation of estimated selection coefficients. In particular, a rescaling factor of 8 means that a single generation in the simulation model corresponds to 8 generations in the real population. Thus, a sweep that would fix in, say, 160 generations in the real population, would correspond to a sweep that fixes in only 20 generations in our rescaled model, therefore requiring a much higher value s. We will show below how this reasoning can be used to map an inferred selection coefficients from the rescaled model onto its corresponding values in the real population.

At *Ace*, our method classified the sweep as an RNM soft sweep with probability 98.8 % to 99.9 % (median 99.6 %), consistent with the known sweep mechanism. The selection coefficient was inferred to be between 1.82 < *s* < 3.35 (median 2.41) across 10 model training replicates. A sweep with *s* = 2.41 in our rescaled simulation model takes on average ~ 42 generations to fixation. Given the rescaling factor of 8, this should correspond to ~ 336 generation in the unscaled population with *N_e_* = 400, 000. Using Wright-Fisher simulations, we estimated that this corresponds to a selection coefficient of s ~ 0.14 in the unscaled population, which is broadly consistent with previous estimates (Karasov et al., 2010). The sweep at *CHKov1* was correctly inferred as a soft sweep from SGV, with all model replicates giving a probability above 99.9 %. The selection coefficient was inferred to be between 15.5 < *s* < 57.1, with a median value of 41.7. While this value may appear very large, it specifies a sweep that on average still takes ~ 22 generations to fixation in our rescaled model, and thus is only about twice as fast as the inferred sweep at *Ace* (thereby providing a nice illustration for how the scaling of selection coefficients becomes far from linear for larger *s*). This should correspond to a sweep taking ~ 176 generations in the unscaled population, which yields s ~ 0.28. Finally, the sweep at *Cyp6g1* was again correctly inferred as a soft sweep, with our method classifying it as SGV sweep with probability 99.1 % to 99.9 % and median 99.9 %. The selection coefficient was inferred to be between 5.16 < *s* < 23.53 (median 7.13). This specifies a sweep that on average takes ~ 29 generations to fixation in our rescaled model, yielding a corresponding sweep duration of ~ 232 generations and a value of s ~ 0.21 in the unscaled population. In summary, the classifications of sweep types by our method are consistent with the know sweep mechanisms at each of the three control loci. The estimated selection coefficients suggest very strong selection, which seems consistent with the fact that all of these sweeps are associated with the evolution of resistance against widely used insecticides.

## Discussion

In this study, we presented a supervised machine learning framework for the inference of sweep parameters from patterns of genetic variation observed around a sweep locus. We demonstrated the performance of our method on models trained for the estimation of selection coefficient and the classification between hard sweeps, SGV soft sweeps, and RNM soft sweeps across a wide range of evolutionary scenarios. We further demonstrated how training data can be customized to adapt the method to new questions, such as an extension to partial sweeps. Our method correctly recovered the sweep types at three loci in *D. melanogaster* where strong selective sweeps of known mechanism have recently occurred. These results suggest that different sweep modes indeed leave distinct signatures in the patterns of surrounding variation that can allow us to infer the strength and type of a sweep with some accuracy.

One critical consideration for any machine learning approach is deciding how to represent the data that is fed into the method (Halevy et al., 2009; Mughal & DeGiorgio, 2019). In our case, we selected a variety of summary statistics evaluated around the sweep locus, which include estimates of the level of nucleotide diversity, the shape of the site-frequency spectrum, and haplotype patterns. Previous approaches have used a similar set of statistics (Schrider & Kern, 2016), while others have suggested alternative representations such as the full site-frequency spectrum (Ronen et al., 2013), haplotype-frequency spectrum (Messer & Neher, 2012), the inferred genealogies (Ralph et al., 2020), or even a picture of the raw genotype alignment in the hope to retain as much original information from the data as possible without any summarization (Flagel et al., 2019). We are not aware of any systematic analysis that has yet tested these representation alternatives against each other under comparable circumstances, so it remains unknown if any of them is consistently more powerful than the others. One important advancement of our method compared to previous approaches is that we systematically varied the window sizes over which summary statistics are estimated. This strategy allows our method to attain power across a wide range of sweep strengths, including very strong sweeps with selection coefficients *s* ≫ 1 (which are not unusual in simulations where evolutionary parameters need to be rescaled for computational feasibility).

We join previous authors in arguing that supervised machine learning can be a powerful strategy for estimating population genetics parameters. Since evolutionary history in nature is hard to know, and there are still few cases of ground-truth knowledge of selective sweeps or demographic history, such methods typically rely on simulated training data. We argue that this family of methods belongs to the transfer learning paradigm, where a model is trained on a source domain of data before being applied to a different but related target domain (Weiss et al., 2016). Importantly, these methods need to address the possibility of negative transfer: if the source and target domains are too dissimilar, results in the target domain might be misleading (Rosenstein et al., 2005). In this work, we considered three selective sweeps in *Drosophila melanogaster* to be labeled data in the target domain and used them as controls in order to test the performance of the method *a posteriori*. Formal transfer learning algorithms are available where performance on the target domain is used to inform the training process in the source domain (Weiss et al., 2016). These algorithms have the advantage of being able to quantify negative transfer as well. We anticipate that they will become more prevalent in the field as the amount of labeled evolutionary data increases and evolutionary simulation software continues to improve. Tools for model interpretation of machine learning output, both model-agnostic (Ribeiro et al., 2016) and specific to neural networks (Olah et al., 2018), are also available and can be used to improve interpretation of results.

We tested the performance of our method under a highly idealized evolutionary model of a panmictic population of constant size. However, the flexibility of the SLiM simulation framework used for generating the training data allows simulations to be tailored for any specific organism and evolutionary scenario. Demographic history, population structure, or any aspect of mating or life history can be easily incorporated in SLiM, which also provides direct support for the growing set of standardized evolutionary models implemented in the stdpopsim library (Adrion et al., 2020). Strategies to make simulations even more realistic could include varying levels of dominance and models of older sweeps sampled some time after fixation, as both factors can affect sweep signatures (Hartfield & Bataillon, 2020; Przeworski, 2002). Training data could also be simulated to incorporate missing data and sequencing error. Even further customization could be achieved by tailoring simulations to the specific locus of interest, given that mutation and recombination landscape variation, background selection, presence of nearby genes, and recurrent sweeps can all affect sweep signatures and interact with each other in ways that are often hard to predict.

The question arises of how much tailoring of simulations to do. Is it ideal to create a highly customized dataset with fixed parameters that are known about the population and locus of interest, or is it better to train the model with a more general parameter distribution? In practical applications, the precise evolutionary parameters of a study population are often fraught with considerable uncertainty. It has therefore been suggested that evolutionary parameters should be tuned to maximize the fit of the used summary statistics to those observed in the real data (Garud et al., 2021). An alternative approach is to model evolutionary parameters as random variables in the training data to account for uncertainty and to allow the the model to learn about a more generalizable distribution of scenarios. For instance, in this work, we have opted to train a single model with a distribution of partial sweep frequencies, rather than training one model per locus with data containing only the specific partial frequency (if known) of the sweep in question. Similarly, if one wants to estimate the selection coefficient of a sweep one presumes to be an SGV soft sweep, it is unknown if it were best to use a method trained on only such sweeps, or if it is better to still include other sweep types in training, as we have done here. Choosing which approach to take likely requires a trade-off: a more general training dataset can be more difficult to train on and have noisier estimates, but it is presumably more robust to overfitting than a highly tailored one with fixed parameters, especially if there is uncertainty about the true parameter values. One systematic way of guiding the choice is to treat this question as a problem of hyperparameter tuning (Abu-Mostafa et al., 2012), choosing the set of simulation parameters that maximizes performance on an independent validation dataset.

Supervised learning is a tool that we believe should be easily available and widely applicable by the field of population genetics. The approach we developed in this work serves that purpose by introducing a forward simulation framework that can be intuitively customized and extended to fit a study organism or locus of interest and then used as training data for a model capable of inferring any given sweep parameter of the simulations. Our models inferred selection coefficient and sweep type, but supervised learning is a general framework and other parameters of evolutionary interest can be inferred from data following the same approach. We thus hope that our framework can contribute to lowering the technical barrier of parameter inference in population genetics.

---

**Supporting Figure 1:**
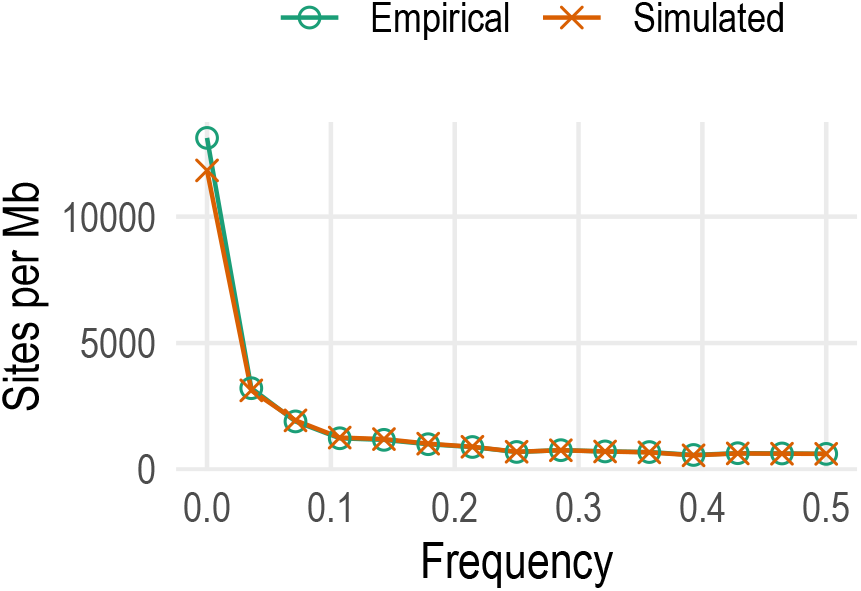
Site-frequency spectrum of neutral simulations and empirical data. The *y*-axis counts the number of segregating sites in one megabase, averaged across the genome for the empirical data and across simulations for the simulated data.

**Supporting Figure 2:**
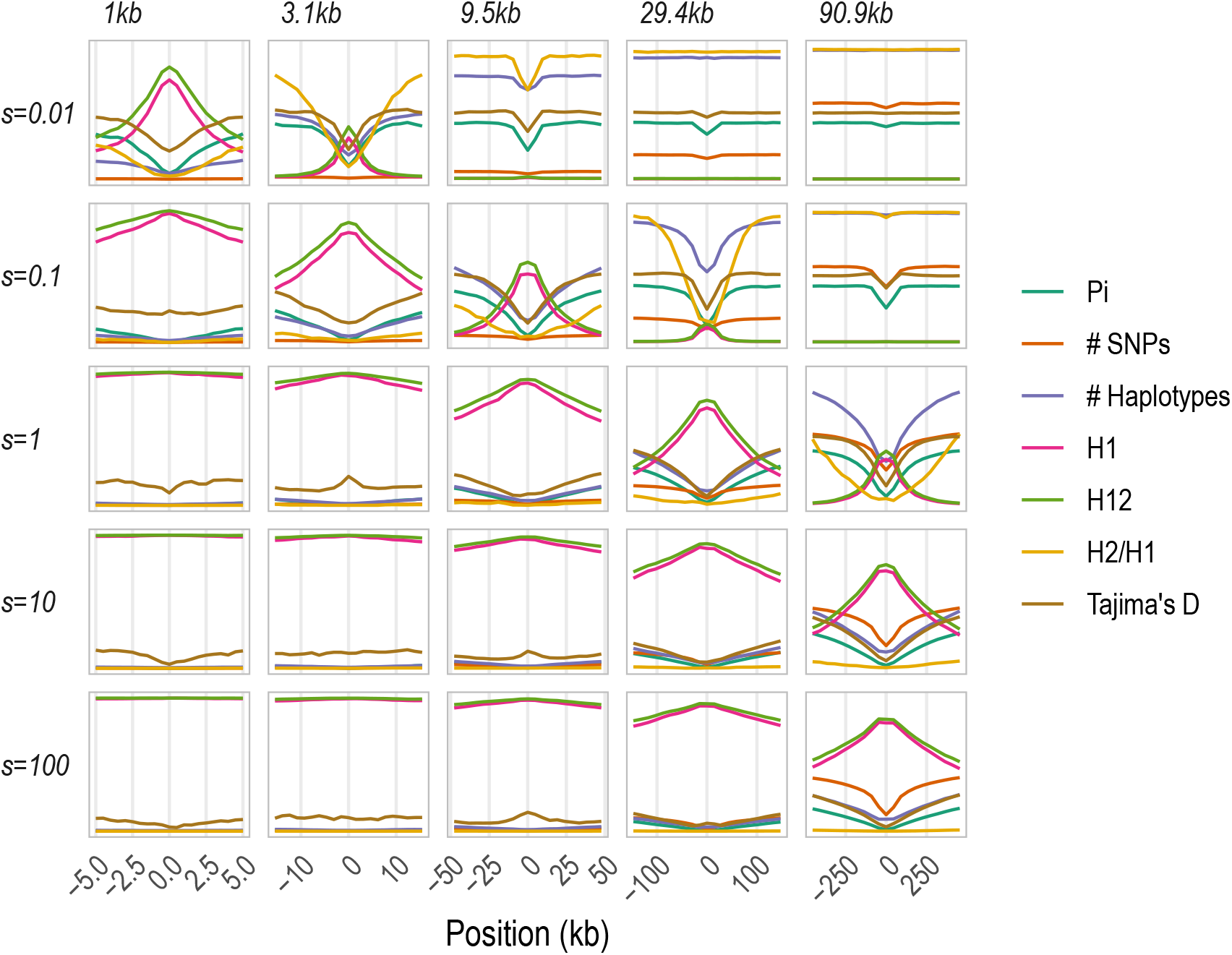
Sweep signatures of hard sweeps averaged over 100 simulations, with five out of 21 subwindow sizes shown for compactness. The *y*-axis shows normalized statistic values as described in the main text.

**Supporting Figure 3:**
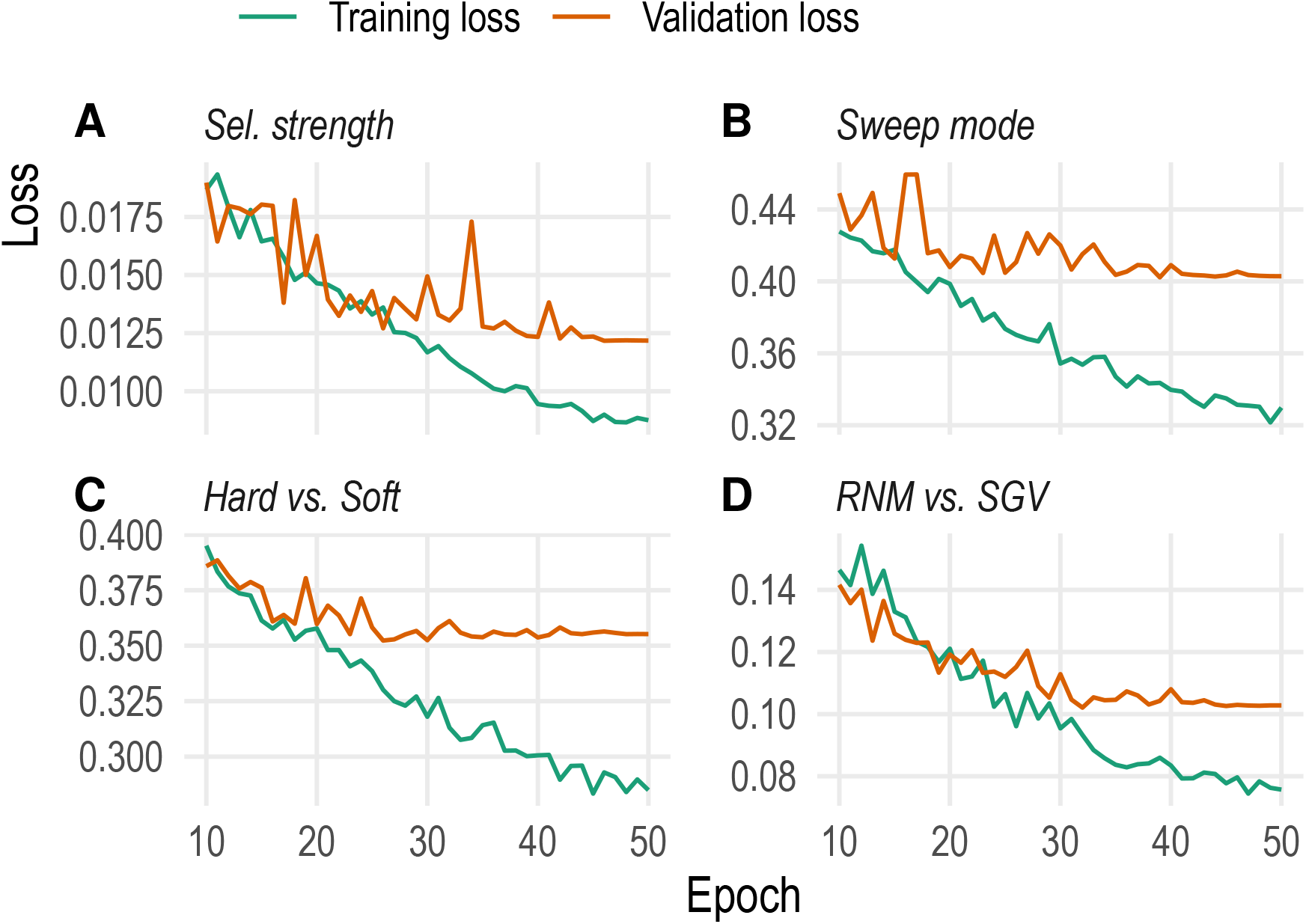
CNN learning curves of the machine learning models.

**Supporting Figure 4:**
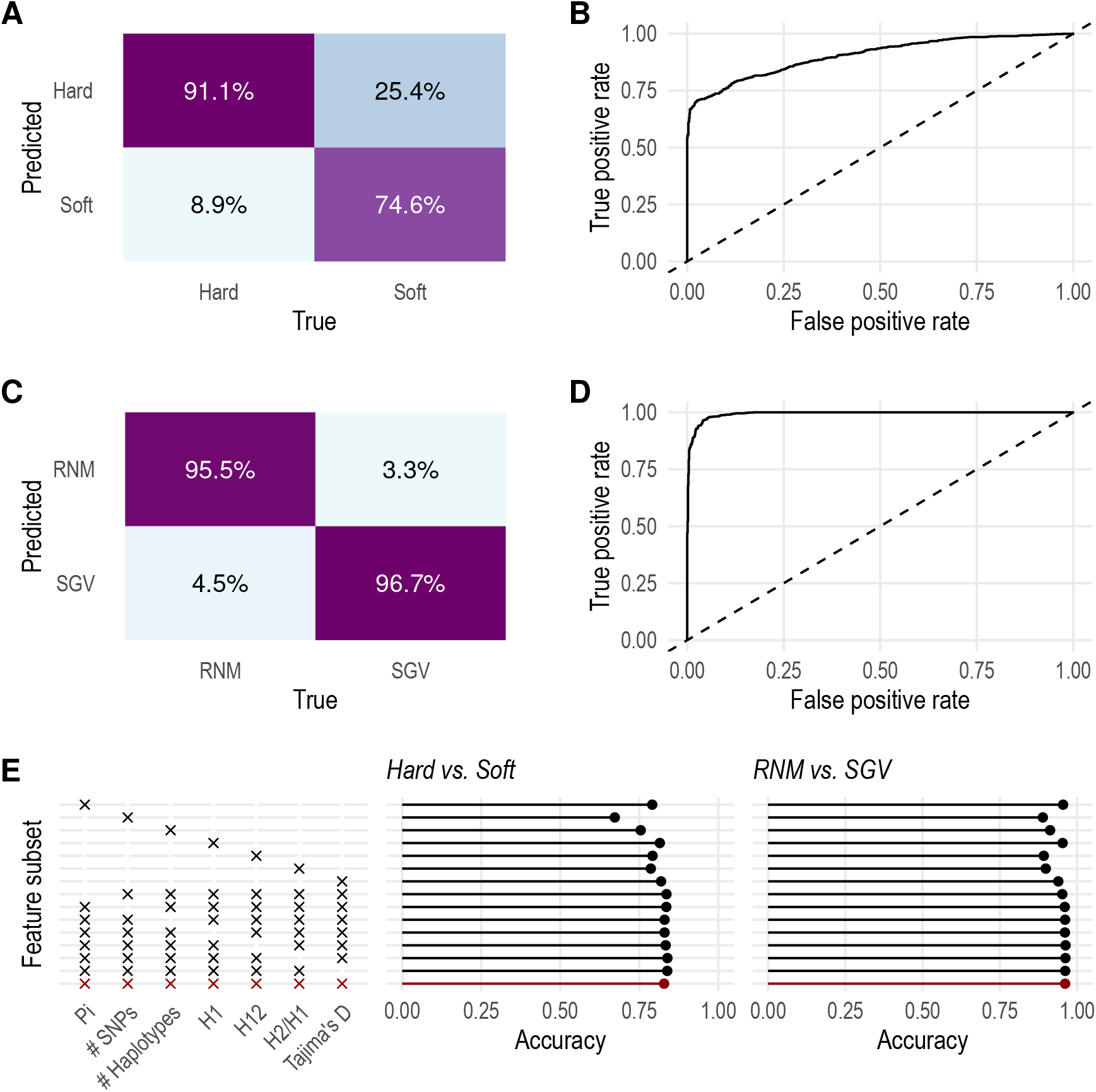
Validation performance of two models of binary classification: A model to distinguish hard from soft sweeps (panels A and B) and a model to distinguish RNM from SGV soft sweeps (panels C and D). (E) Performance validation by subset of summary statistic. Each row is a separate scenario with only the statistics marked with an “x” on the left panel included in training. The baseline scenario with all statistics is highlighted in red.

**Supporting Figure 5:**
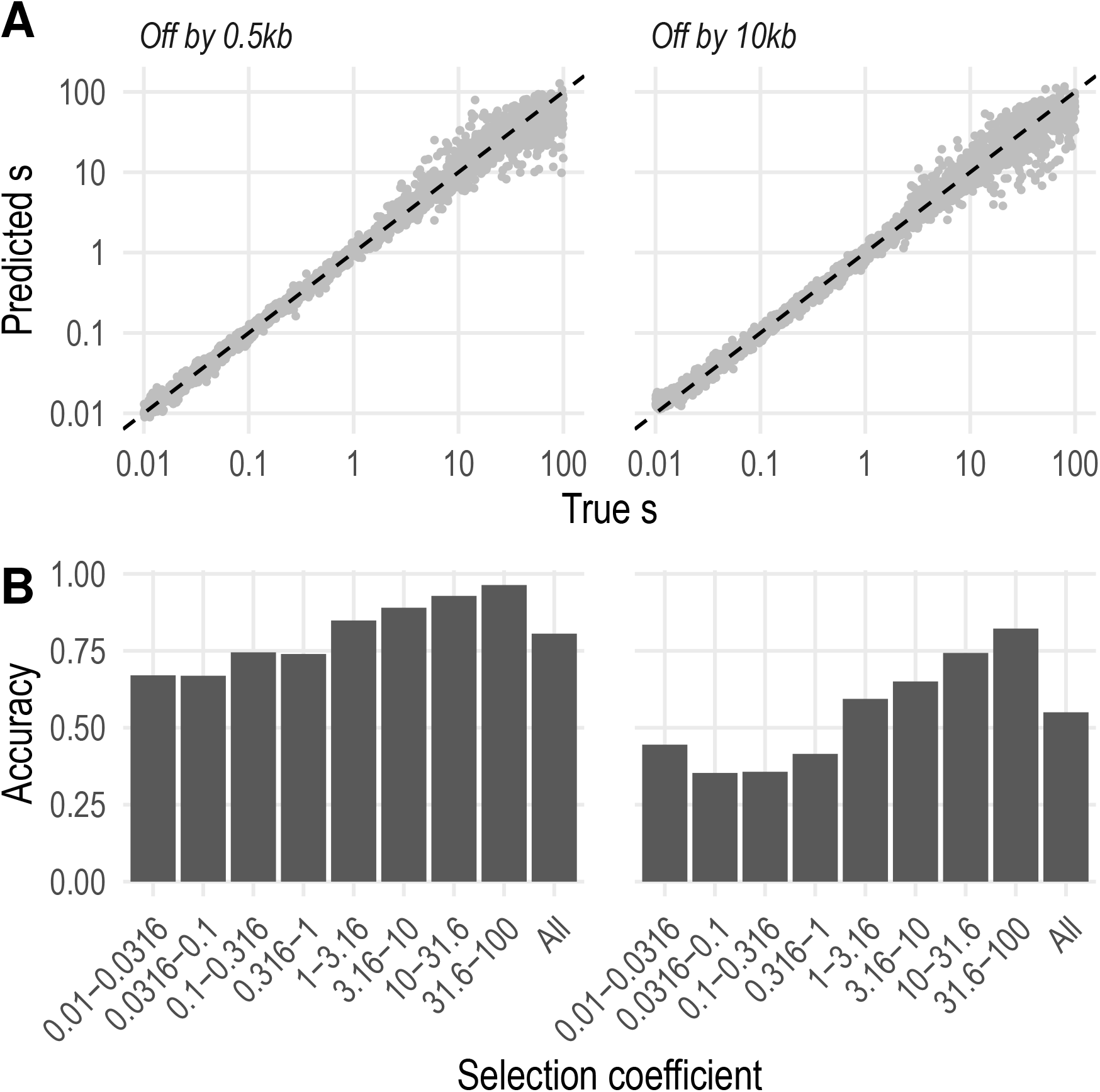
Performance of machine learning models when applied to sweeps not in the middle of the window. (A) Selection coefficient estimation. (B) Accuracy of sweep mode inference.

**Supporting Figure 6:**
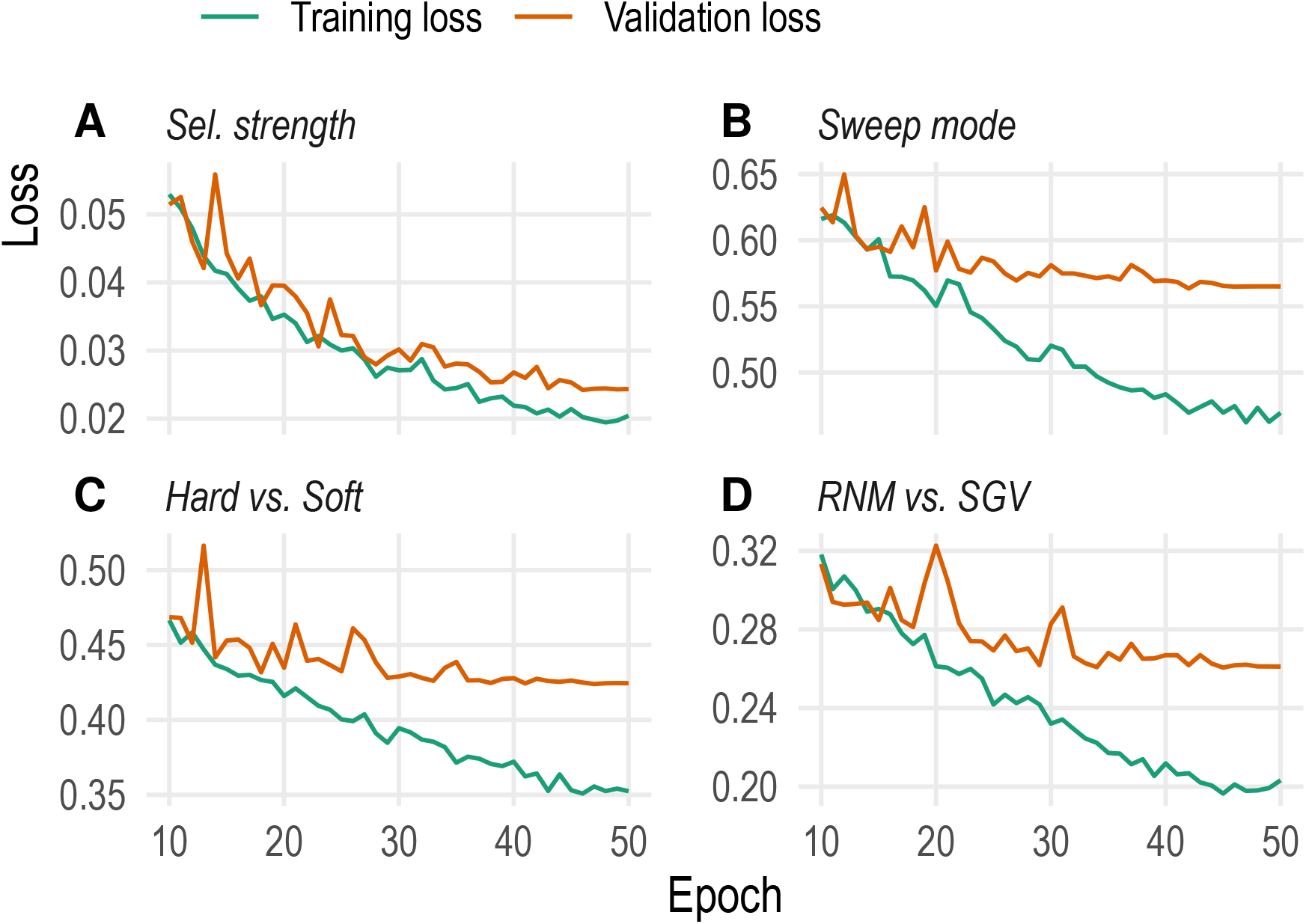
CNN learning curves of the machine learning models trained on partial sweeps.

**Supporting Figure 7:**
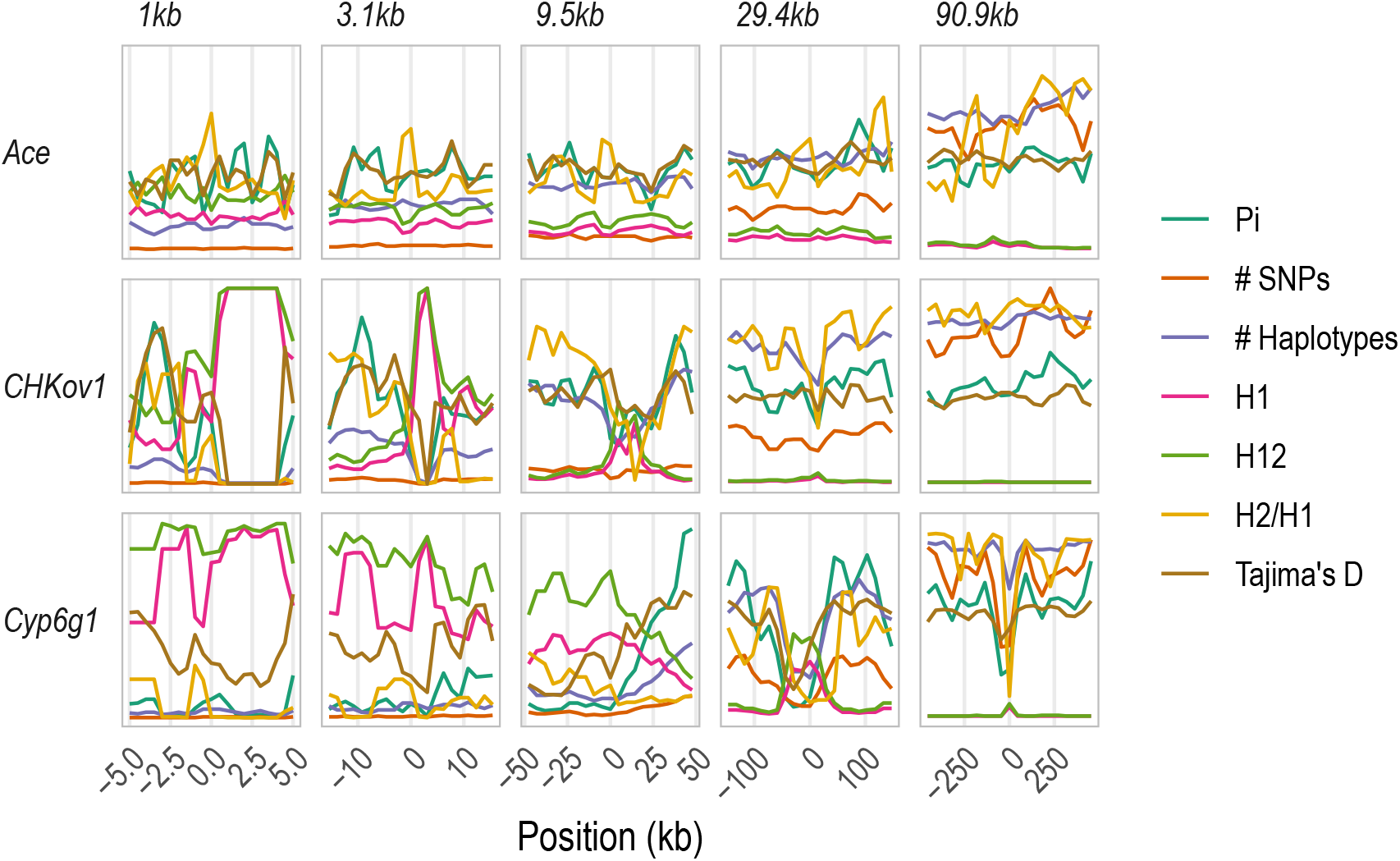
Signatures of selective sweeps at three control loci in *Drosophila melanogaster*. Five subwindow sizes out of 21 are shown for compactness. *CHKov1* and *Cyp6g1* have valleys of heterozygosity at more than one subwindow resolution, while *Ace* displays a large valley of heterozygosity extending beyond the boundaries of the analyzed region. This could be because the evolutionary scenario at *Ace* was more complex than a single sweep, possibly involving several sweeps in the surrounding region.

**Supporting Figure 8:**
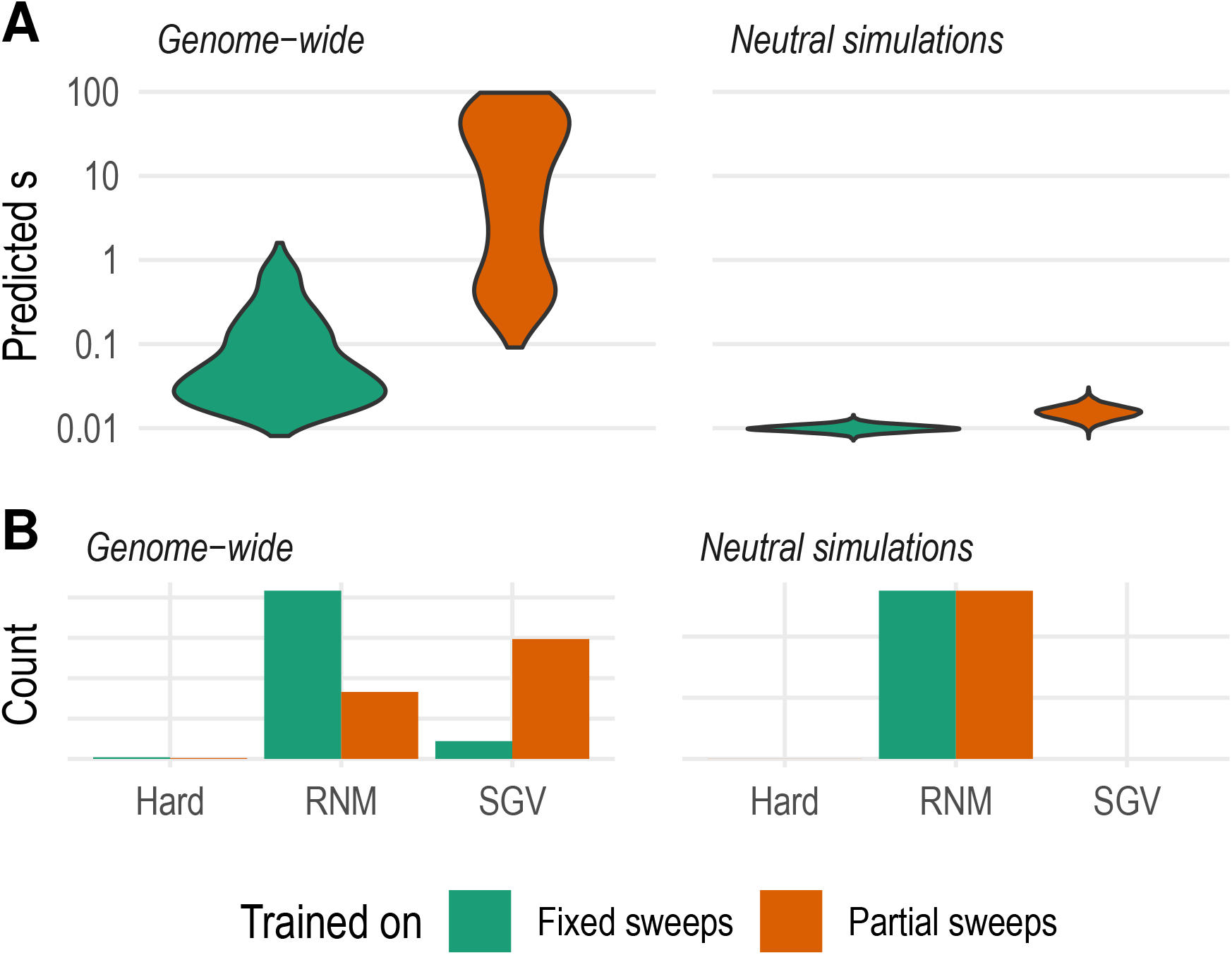
The method always assumes there is a selective sweep at the center of the focal genomic region. We applied the trained models to simulated and empirical regions free of sweeps to test how it performs under violation of this central assumption. Simulation validations were done with a dataset of 5000 neutral coalescent simulations. Empirical validations were done with windows taken from the 2L, 2R, 3L, and 3R chromosomes of the DGRP2 dataset. (A) Selection strength inference. Both models detect very narrow ranges of “selection strength” for neutral simulated windows with modes near the smallest training value of 0.01, the expected guess for a region with no true selection. The empirical genome-wide estimates have wider distributions. In both cases, the model trained on partial sweeps infers a higher selection strength than the one trained on fixed sweeps. The weakest sweeps are the only data in the training dataset of the first model that resemble neutral and empirical windows, but those regions might resemble incomplete sweeps with higher selection strength in the second model’s training dataset. It’s worth noting that genome-wide windows of *D. melanogaster* are not seen as equivalent to neutral simulated windows, presumably because the *D. melanogaster* genome does not show signatures of classic neutrality due to density of positive and background selection (Andolfatto, 2007; Comeron, 2014; Li & Stephan, 2006). (B) Sweep mode inference. The model trained on fixed sweeps classifies both neutral simulated windows and the genome-wide data as soft sweeps from recurrent *de novo* mutation, as they are are the sweeps with the most genetic diversity that it knows about. The model trained on partial sweeps, in contrast, finds RNM sweeps for simulated windows but splits its genome-wide results between soft sweeps from RNM and SGV. Because samples of partial sweeps have higher genetic diversity than ones of fixed sweeps, the model has learned to differentiate between the modes of soft sweep even in the face of high heterozygosity. All in all, the parameter estimates made by our method are explainable even under violation of the central assumption of a selective sweep in the center of the window.

**Supporting Table 1:** Bounds used to normalize raw values of summary statistics.

| Statistic | Lower bound | Upper bound |
| --- | --- | --- |
| SNP count | 0 | 4216 |
| $\pi$ | 0.00 | 0.0106 |
| Tajima's $D$ | -3.00 | 3.00 |
| Haplotype count | 0 | 205 |
| $H_1$ | 0.00 | 1.00 |
| $H_{12}$ | 0.00 | 1.00 |
| $H_2/H_1$ | 0.00 | 1.00 |

**Supporting Table 2:** Subwindow sizes used to capture different resolutions of data.

| Subwindow size (kb) | Covered (kb) |
| --- | --- |
| 1 | 11 |
| 1.253 | 13.783 |
| 1.570 | 17.270 |
| 1.967 | 21.637 |
| 2.464 | 27.104 |
| 3.088 | 33.968 |
| 3.869 | 42.559 |
| 4.847 | 53.317 |
| 6.074 | 66.814 |
| 7.610 | 83.710 |
| 9.535 | 104.885 |
| 11.946 | 131.406 |
| 14.968 | 164.648 |
| 18.754 | 206.294 |
| 23.498 | 258.478 |
| 29.441 | 323.851 |
| 36.888 | 405.768 |
| 46.219 | 508.409 |
| 57.909 | 636.999 |
| 72.557 | 798.127 |
| 90.909 | 999.999 |

**Supporting Table 3:**
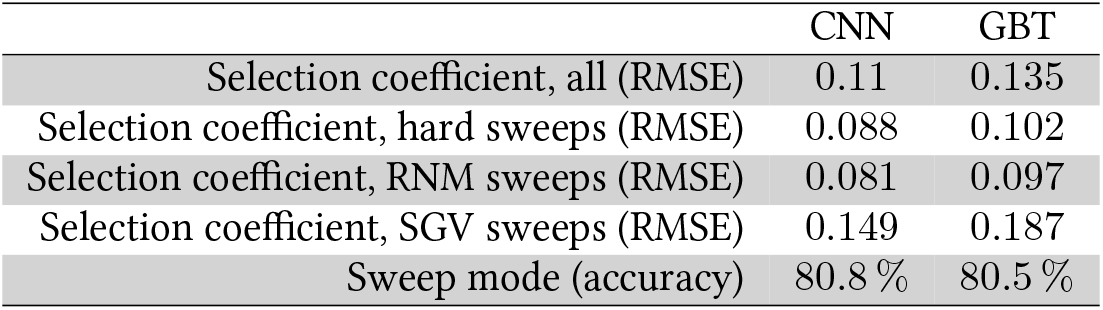
Validation of convolutional neural networks (CNN) compared to Gradient-boosted trees (GBT). RMSE=Root mean squared error.

|  | CNN | GBT |
| --- | --- | --- |
| Selection coefficient, all (RMSE) | 0.11 | 0.135 |
| Selection coefficient, hard sweeps (RMSE) | 0.088 | 0.102 |
| Selection coefficient, RNM sweeps (RMSE) | 0.081 | 0.097 |
| Selection coefficient, SGV sweeps (RMSE) | 0.149 | 0.187 |
| Sweep mode (accuracy) | 80.8 % | 80.5 % |

